# Quantitative measures of topographic and divergent/convergent connectivity in diffusion MRI of the human cerebral cortex

**DOI:** 10.1101/2022.12.25.521904

**Authors:** Liang Shi, Alexander Woodward, Jun Igarashi

**Author notes:** Corresponding author at: Jun Igarashi, High Performance Artificial Intelligence Systems Research Team, Center for Computational Science, RIKEN, Japan.

## Abstract

Spatial features of connections, such as topography and divergence/convergence, reflect the information-processing mechanisms crucial for understanding and modeling the brain. However, this has not yet been comprehensively investigated. Using diffusion Magnetic Resonance Imaging (dMRI) data, we developed a topographic factor (TF) and divergence/convergence factor (DC) to quantitatively explore the spatial connectivity patterns of one entire hemisphere of the human cerebral cortex. In the analysis, the early sensory areas, which are located far away from the center of the cerebral cortex, exhibited high topographic connectivity. In contrast, the limbic system, which is located proximal to the center, showed high divergence/convergence in two types of connectivity: one region to another region at the region-to-region level, and one region to all other regions at the global level. Topography had anti-correlation with divergence/convergence over the cerebral cortex, and the two types of divergence/convergence displayed positive correlation with each other. These results suggest that topographic and divergent/convergent connectivity are spatially organized with respect to cytoarchitecture over the cerebral cortex to optimize energy efficiency and information transfer performance in the cerebral cortex.

## Introduction

The brain is spatially organized, as observed in the structure of connections (y Cajal, 1899), regions (Brodmann, 1909; Glasser et al., 2016), layers, columns (Mountcastle, 1997), and topographic organization (Tanaka, 1997; Hagler Jr and Sereno, 2006; Silver and Kastner, 2009; Harvey et al., 2013; Gu et al., 2018; O’Rawe and Leung, 2020). These features have emerged during the evolutionary process for efficient information processing while conserving energy (Bullmore and Sporns, 2012). It is crucial to understand their organization in order to decipher the mechanisms of information processing and neuronal disorders. In particular, brain connectivity is a cardinal feature of the spatial organization and has been extensively studied (y Cajal, 1899; Oh et al., 2014).

Two core features of spatial connectivity have been observed in the brain, namely topographic and divergent/convergent connectivity. Topographic connectivity is the preservation of the spatial relationship between neurons in source and target regions, which is marked in early sensory areas, such as the retinotopy in the early visual cortex (Dumoulin and Wandell, 2008; Wang et al., 2015), tonotopy in the early auditory cortex (Leyton and Sherrington, 1917; Romani et al., 1982), and somatotopy in the early sensory-motor cortex (Harvey, 1909). Essentially, this pattern is recognizable over the entire brain (Thivierge and Marcus, 2007). The functional role of topographic connectivity is considered to be for the processing of local sensory features and even for abstract processing such as analogical reasoning (Thivierge and Marcus, 2007). Divergent and convergent connectivity scatters connections from source to target, and gather connections from target to source, respectively. This pattern is found abundantly in the hippocampal and parahippocampal areas (Suzuki and Amaral, 1994; Wellman and Rockland, 1997) and also widely observable over the entire brain. The major functional role of divergent/convergent connectivity is considered to be for integrating and broadcasting information for emotion, decision making, multimodal sensory information, memory retrieval/encoding, and navigation (Sporns et al., 2002).

These two types of connectivities are fundamental components of the wiring in the brain. However, how they are organized over the cerebral cortex from the viewpoint of anatomical classification (e.g. Terminologia Anatomica(FCAT, 1998)), cytoarchitectonic features, and functional role in information processing has not been comprehensively investigated. We therefore used diffusion Magnetic Resonance Imaging (dMRI) in this study to allow us to analyze the entire human brain quantitatively. dMRI is a widely used non-invasive method that reflects the orientation of axonal fibers by measuring the diffusive motion of water *in-vivo* (Tournier et al., 2011). A number of studies have used dMRI data to elucidate topological features (Fornito et al., 2016), and disease-related connectivity in patients with Alzheimer’s disease (Ye et al., 2019) and Parkinson’s disease (Barbagallo et al., 2017).

In this study, we investigated the topographic and divergent/convergent connectivity in one hemisphere of the human cerebral cortex at the dMRI voxel resolution. First, to confirm topographic features in the cortex, we examined the topographic connectivity between the primary motor and sensory cortex, which are known to have clear topographic organization. We further extended our analysis to the early visual cortex and the entirety of cortical areas. Next, we analyzed the divergent/convergent connectivity at two spatial scales: (a) region-to-region and (b) one region to all others regions in the entire cortex (global). Finally, we investigated the relationship between the topography and divergent/convergent connectivity. We found high topographic connectivity of the early sensory cortices in contrast to high divergent/convergent connectivity of the limbic system. These results suggest that topographic and divergent/convergent connectivity may be spatially organized according to the varying type of regions in the cortex, optimizing between connection cost and functional roles. This is the first comprehensive and quantitative study on topography and divergence/convergence measures over the entire cortex using dMRI.

## Materials and Methods

### Subjects and imaging data

T1-weighted (T1w) structural MRI and dMRI data were obtained from the Human Connectome Project (HCP) database (Essen et al., 2013). Owing to limited availability of computational resources, we randomly selected 11 subjects (5 male, 6 female) from 100 non-related subjects released in HCP 1200 subject to exclude confounding effects. The IDs of these subjects were 100307, 105115, 110411, 111312, 115320, 120111, 123117, 127933, 147737, 163129, 280739 and all of them were healthy, young people between the ages of 25 and 35 years (31.2 ± 2.5). Preprocessed T1w MRI data (0.7×0.7×0.7*mm*) and 3T dMRI data (1.25×1.25×1.25*mm*) from all subjects were used for analysis. The MRI acquisition and preprocessing protocols are described in (Sotiropoulos et al., 2013; Glasser et al., 2013).

### Structural connectivity generation

There are two main steps in generating a voxel-wise connectivity matrix: individual parcellation generation in the diffusion space and tractography reconstruction. For the first step, according to the HCP’s multimodal cortical parcellation (HCP_MMP1.0) (Glasser et al., 2016), the cerebral cortex of each subject was parcellated into 360 regions of interest (ROIs) by co-registering each individual T1w image with the common surface-based template, fsaverage template mapped with the HCP_MMP1.0 atlas’ annotation file^1^ using the FreeSurfer software package (Fischl, 2012). From this an individual parcellation in T1w space was acquired. A linear image registration tool, FLIRT, was then used to calculate a transformation matrix, *T*, that transforms images from the T1w space to the diffusion space for each subject (Jenkinson and Smith, 2001; Jenkinson et al., 2002). Finally, individual parcellation in the diffusion space was obtained using *T*.

The next step was to generate tractography. Diffusion data were processed following the MRtrix3 guidelines for generating structural connectivity (Tournier et al., 2019). In summary, an orientation distribution function at each voxel was estimated by multi-shell multi-tissue constrained spherical deconvolution method (Jeurissen et al., 2014). Anatomically-constrained tractography with back-tracking was performed since anatomical information can significantly alleviate false positives (as observed when comparing to neuronal tracer data as ground truth (Smith et al., 2012; Aydogan et al., 2018)), and the back-tracking method better detects plausible pathways within narrow gyral projections (Smith et al., 2012). Each voxel in the cerebral cortex was labelled a unique integer value and considered as one seed to reconstruct tractography using probabilistic tracking (iFOD2, maximum length: 250*mm*, FA cutoff value: 0.05) (Tournier et al., 2010). Compared with deterministic tractography, probabilistic techniques can better detect non-dominant fiber pathways (Behrens et al., 2007). The tracking process was stopped after generating 5000 streamlines from each seed voxel, or until 5000000 tracking attempts were attempted. For reducing bias, connectivity from the seed voxel was discarded when less than 5000 streamlines were generated. Finally, we built a voxel-wise connectivity matrix of the cerebral cortex and did this for each participant (Fig. 1a). To obtain a sparse whole-brain connectivity matrix, all values in the connectivity matrix less than 10 were zeroed, and all connections to the target region were discarded if the number of voxels with connections in the target region was less than 10. Additionally, since we focused on inter-regional connections within one hemisphere, only the 180 ROIs of the left hemisphere in the cerebral cortex were used.

**Figure 1:**
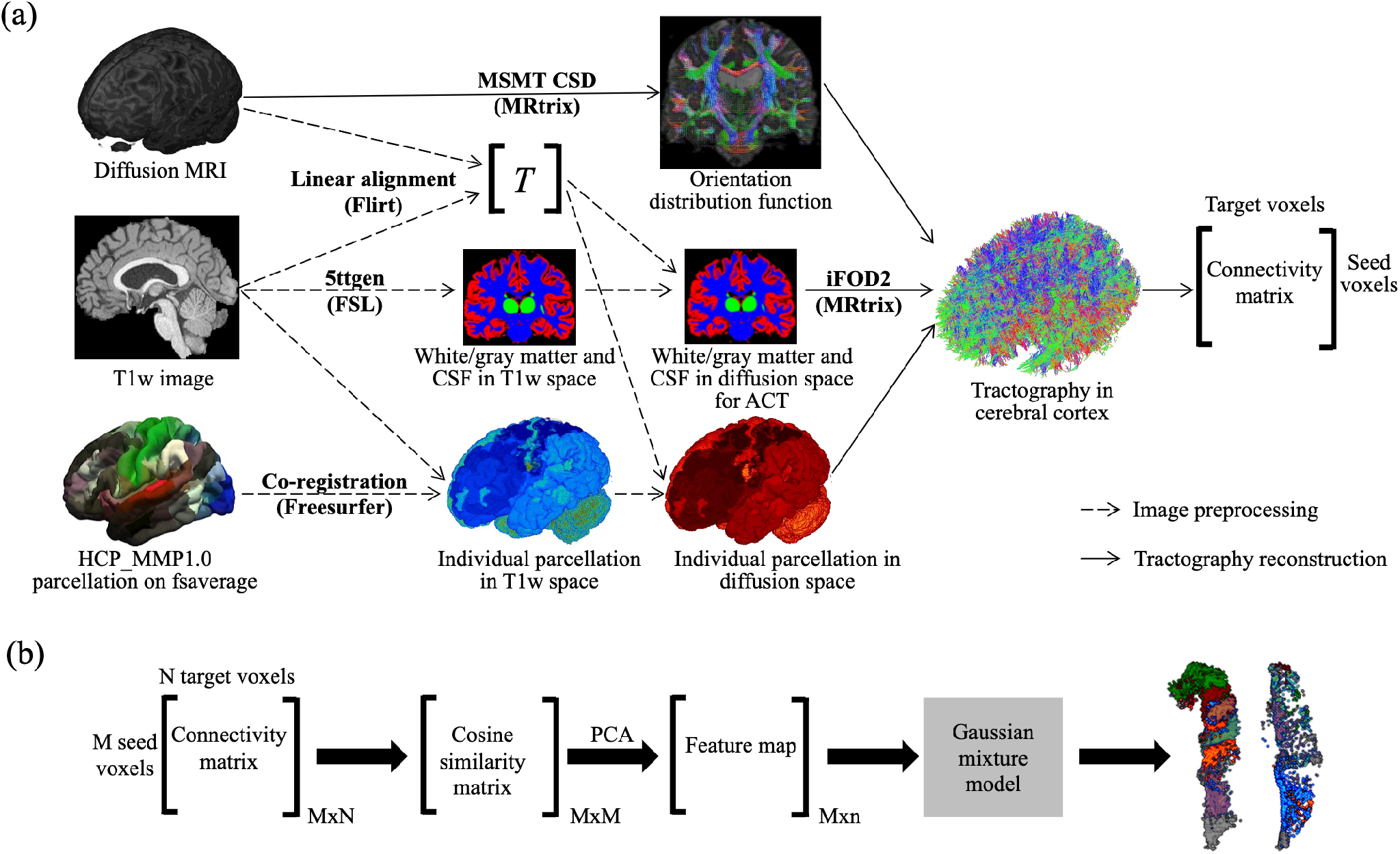
(a) Overall procedure to generate voxel-wise connectivity matrix. MSMT CSD: multi-shell multi-tissue constrained spherical deconvolution; ACT: anatomically-constrained tractography. (b) Flowchart for parcellating M1 and S1 into subregions based on the connectivity matrix. (Brain image examples were from subject 100307.)

### M1 and S1 segmentation

To segment M1 and S1 into several subregions, we used the ROIs in the Montreal Neurological Institute (MNI) space as standard parcellation for subsequent steps^2^. The rigid, affine transformation, and symmetric image normalization method (SyN) in ANTs (Avants et al., 2009) were used to register ROIs from the individual to the MNI space, and a deformation field, *ϕ*, between the two spaces was generated. Individual ROIs were resampled so that their voxel count was the same as in the standard parcellation, and then each individual voxel was transformed to MNI space using *ϕ*. Next, we matched voxels between the transformed individual ROI and standard parcellation using an optimized assignment method (lap-solver package^3^). The group average connectivity matrix between M1 and S1 was then acquired. With this connectivity matrix we used cosine distance to measure the connectivity similarity of voxels, and principle component analysis (PCA) for dimensionality reduction (*explained variance ratio*>0.955). A feature map was then obtained and the Gaussian Mixture Model (GMM) clustering algorithm (Pedregosa et al., 2011) was used to generate subregions (Fig. 1b). The Bayesian information criterion (BIC) gradient was used to determine the best cluster number.

### Region characterization

Ten features were selected to characterize paired regions: volume ratio, surface area ratio, average aspect ratio, ratio of region aspect ratios, average sphericity, ratio of region sphericities, Euclidean distance, minimum volume, minimum surface area, and average distance from the center of the two hemispheres of the cerebral cortex. Minimum volume, minimum surface area, volume ratio, and surface area ratio (larger over smaller one) were used to measure region size. Aspect ratio is the ratio of the range of the projections of a region’s voxel’s coordinates onto its first and second main PCA directions. Sphericity is a measure of how closely the region resembles a sphere. Both aspect ratio and sphericity can measure the shape of a region. Sphericity is defined as:

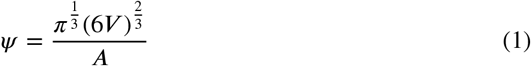

where *V* and *A* are the volume and surface area of the region, respectively. Euclidean distance is the distance between the center coordinates of paired regions. The ratio of region aspect ratios and sphericities were used to evaluate the relative shape relationship between paired regions. The average distance from the center indicates how close to the center of the cerebral cortex the pair is. The center was defined as the average coordinate of all parcellation voxels of the two hemispheres of the cerebral cortex, and Fig. 2 shows this center and the most proximal/distal regions in the cerebral cortex of subject 100307. Additionally, five features were used to characterize each individual region: volume, surface area, aspect ratio, sphericity, and distance from the center of the cerebral cortex. We performed z-score transformation for all features, and evaluated their variance inflation factors (<5) to avoid multicollinearity before performing multiple linear regression analysis.

**Figure 2:**
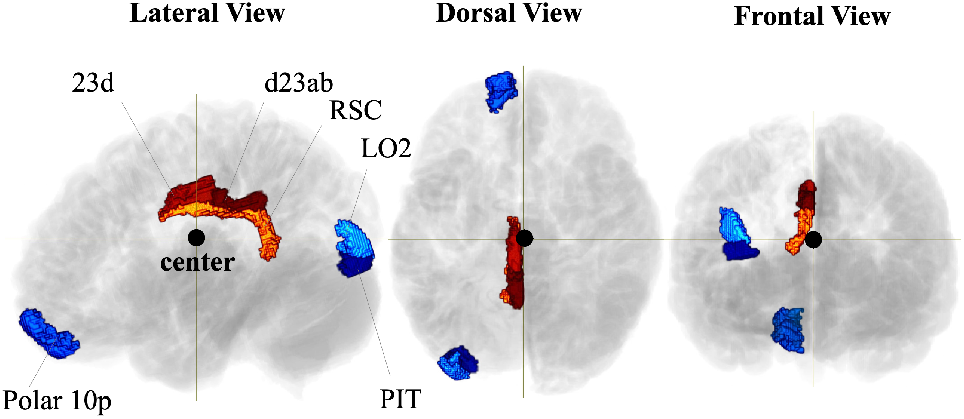
The center of entire cerebral cortex (black circle point). These regions with hot and cool colors represent the regions most proximal and distal to the center, respectively. (Subject: 100307)

### Measure of topographic connectivity pattern

To characterize the topographic connectivity pattern between paired regions, we defined the following three terminologies: source region, target region, and mapping coordinate (Fig. 3). In the source region, technically only a few voxels, named seed voxels, directly connect to target voxels in the target region. The mean coordinate of these target voxels tracked from each seed voxel, using the streamline number as a weight, is calculated to generate a mapping coordinate:

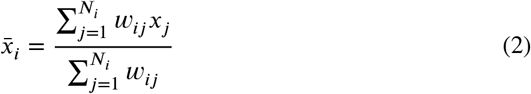

where *i* and *j* are indices of the seed and target voxels, respectively; *x*_*j*_ is the coordinate of the target voxel; *w*_*ij*_ is the number of streamlines from seed voxel *i* to target voxel *j*; 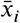 is the mapping coordinate tracked from the seed voxel *i*; and *N*_*i*_ is the number of target voxels tracked from the seed voxel *i*. Topographic connectivity is estimated by performing Procrustes analysis between the coordinates of the seed voxels and mapping coordinates. Procrustes analysis measures similarity between two data sets by calculating the pointwise differences after standardization and optimal linear transformation, including scaling, rotation, and translation (Gower, 1975; Krzanowski, 2000). The topographic factor (TF) is defined as:

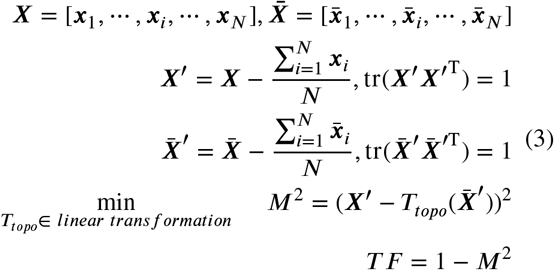

where ***x***_*i*_ and 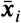 are the coordinates of seed voxel *i* and mapping coordinate, respectively, tr is the trace of matrix. The linear transformation *T*_*topo*_ is applied to 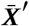 to minimize *M*^2^, and *N* denotes the number of seed voxels within each ROI. TF is in the range of 0 to 1, where 1 represents a perfect topographic connectivity pattern.

**Figure 3:**
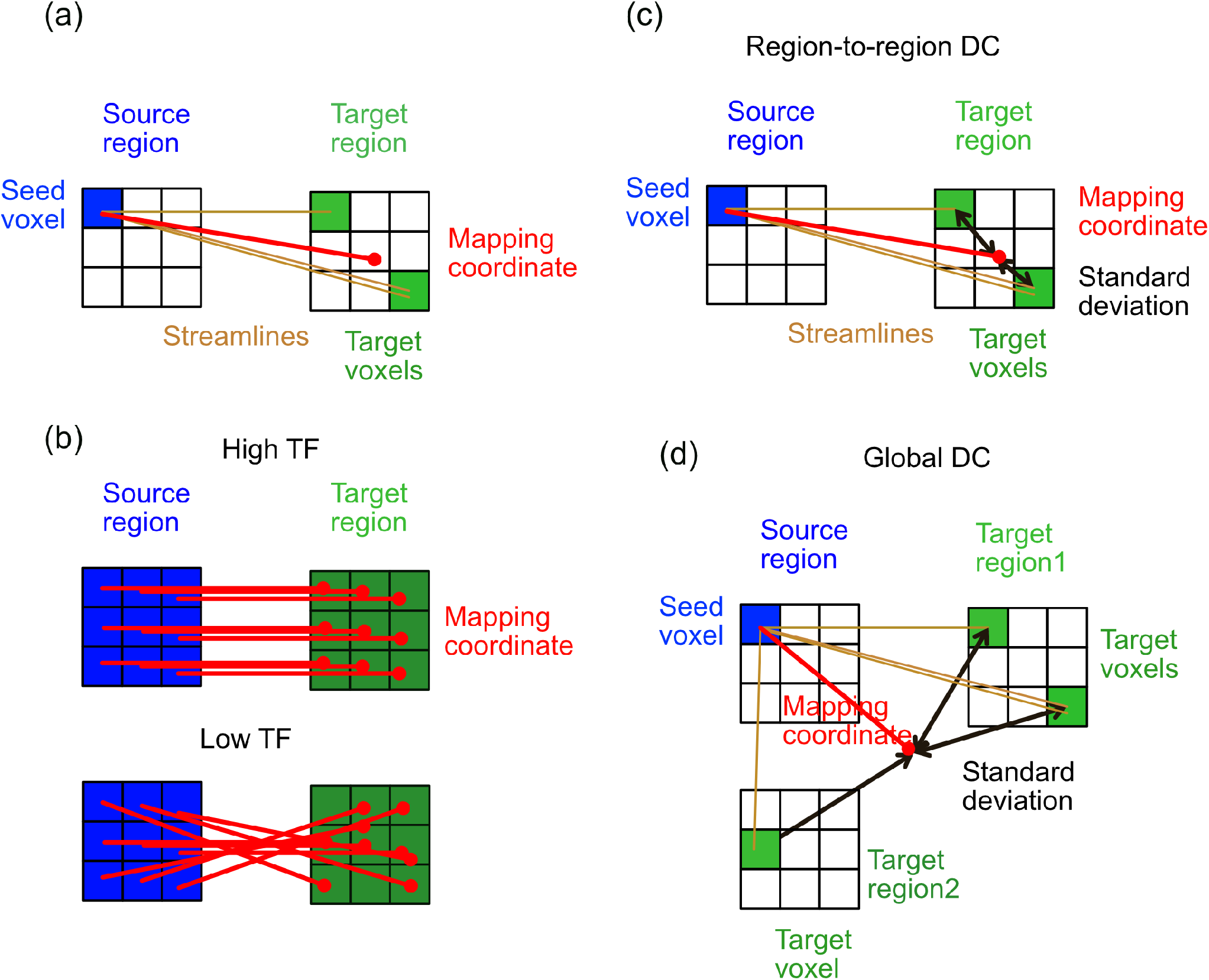
Simplified examples to clarify the definition of TF and DCs. (a) Illustration of defined terminology. The bold red lines represent connections between seed voxels (blue squares) and mapping coordinates (red dots, defined by Eq. (2)). (b) Example of connectivity patterns with high TF and low TF (Eq. (3)). (c) Definition of region-to-region DC. For one seed voxel, the region-to-region DC is the standard deviation of target voxel coordinates from a mapping coordinate, weighted by the number of streamlines within one target region. The final value is the average over all seed voxels in the source region (Eq. (4)). (d) Definition of global DC. For one seed voxel, the global DC is defined as the standard deviation of target voxel coordinates from a mapping coordinate, weighted by the number of streamlines over all target regions. The final value is the average over all seed voxels in the source region (Eq. (5)).

### Measure of divergent/convergent connectivity pattern

The weighted standard deviation of the target voxels tracked from each seed voxel measures the divergent/convergent connectivity pattern, defined as the divergent/convergent factor (DC):

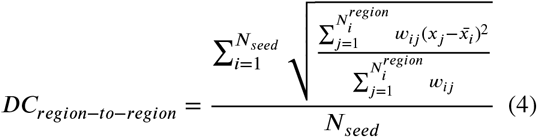

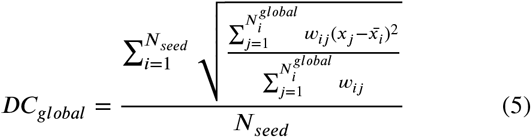

Where *x*_*j*_ and 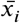 denote the target voxel coordinate and mapping coordinate tracked from seed voxel *i* (defined by Eq. (2)), respectively; *w*_*ij*_ is the number of streamlines from seed voxel *i* to target voxel *j*; *N*_*seed*_ is seed voxel number for each source region; 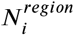 is the target voxel number within each target region tracked from seed voxel *i*; 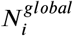 is the target voxel number within one hemisphere of whole cerebral cortex tracked from seed voxel *i*. At the region-to-region level, only those target voxels within one target region were considered. While at the level of the entire hemisphere of the cerebral cortex (global level), we consider all target voxels in all target regions in one hemisphere of the cerebral cortex. Besides, we calculated two values of region-to-region DC for each region pair by swapping the roles of source and target and then used the larger value as the region-to-region DC to give the amount of divergent/convergent connectivity.

## Results

### Topographic connectivity between M1 and S1

Topographic connectivity has been clearly observed in the sensory cortices of the brain (visual, auditory and sensorimotor areas) (Tootell et al., 1988; O’Leary et al., 1999; Weisz et al., 2004; Thivierge and Marcus, 2007; Glasser et al., 2016). If dMRI can be applied to analyze topographic connectivity, it would allow us to comprehensively investigate it over the entire cerebral cortex. First, we confirmed whether it was possible to extract topographic connectivity patterns using dMRI between the M1 and S1 brain regions. Here, topographic organization in the human cerebral cortex has been observed through electrical stimulation (Penfield and Boldrey, 1937), in the connectome of the mouse cerebral cortex (Zingg et al., 2014), and in the functional connectivity of the human and monkey cerebral cortex (Thomas et al., 2021). In the current study, we investigated the degree of topographic connectivity, defined as a topographic factor (TF), which is calculated as a similarity between seed voxel coordinates in the source region and mapping coordinates in the target region using the Procrustes analysis (Eq. (3)). The TFs for M1 to S1 (0.884±0.020, n=11, mean of permutation data by random rematching of seed voxels and mapping coordinates: 0.0007; Mann-Whitney U-test, *P* <0.001) and for S1 to M1 (0.828 ± 0.042, n=11, mean of permutation data: 0.001; Mann-Whitney U-test, *P* <0.001) were both high, indicating strong topographic connectivity between them.

Next, we checked topographic connectivity by visually inspecting the spatial distribution of dMRI connections. We divided the source region into five sub-regions along the longitudinal axis determined by principal component analysis (PCA), and visualized the connection density from each subregion to the target region (Fig. 4a). The peaks of connectivity density in the target region were aligned in the same order as the source subregions along the longitudinal axis for both M1 to S1 and S1 to M1. Note that we examined both directions of streamlines between a pair of regions given non-directionality of dMRI. These results demonstrate that high TF using Procrustes analysis can reflect real topographic connectivity.

**Figure 4:**
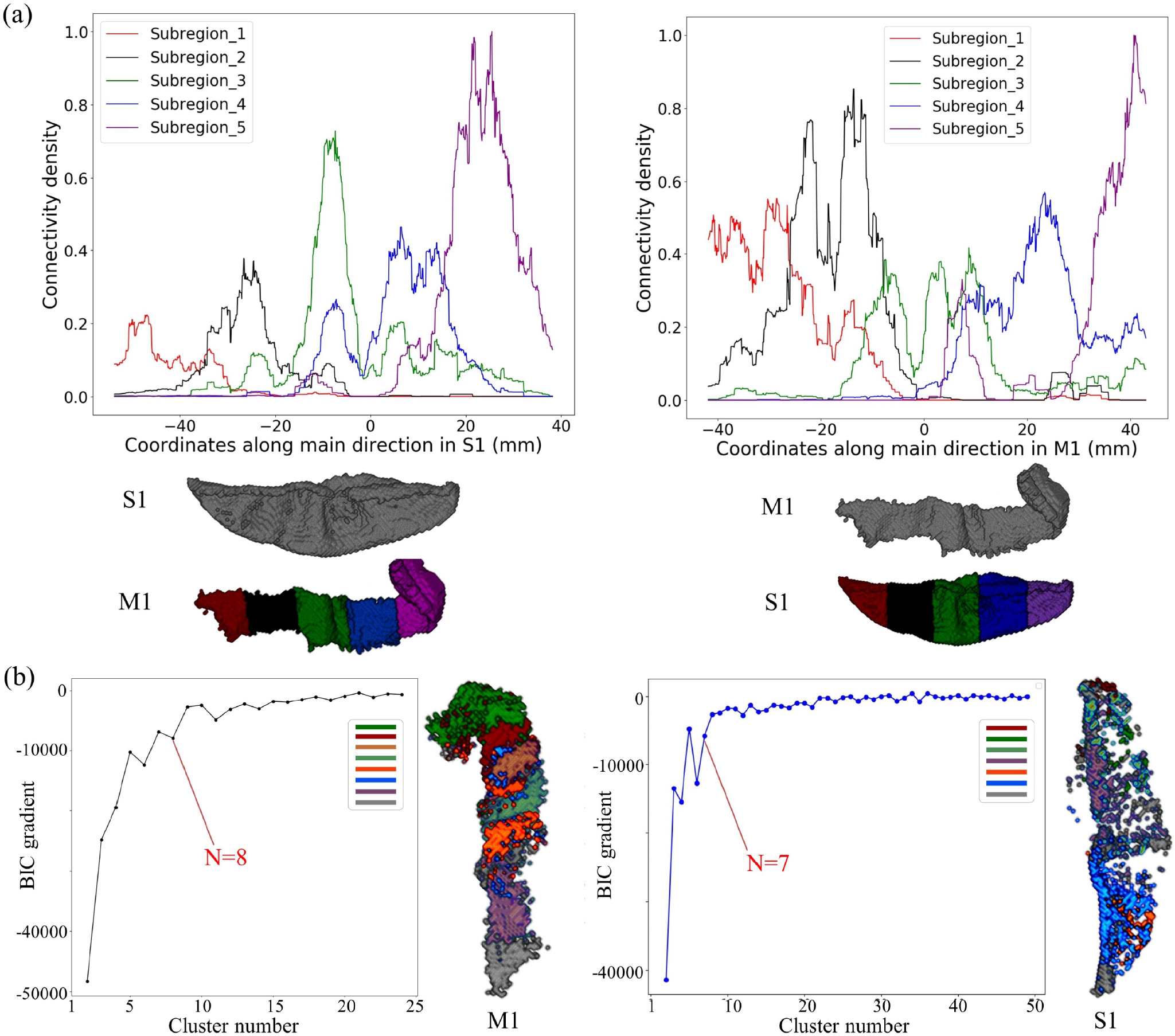
Connectivity pattern analysis between M1 and S1. (a) Connectivity density between each sub-region of M1/S1 and the target region S1/M1, where each color represents one sub-region. (b) BIC gradient and clustering results for M1 (left) and S1 (right).

In the somatotopic maps of M1 and S1, body parts are mapped on the cortical surface along the longitudinal axis (Glasser et al., 2016). To address whether such an alignment of the neural population can be captured from dMRI, we performed clustering analysis by the Gaussian Mixture Model (GMM) algorithm (Pedregosa et al., 2011) and using the connectivity matrix between M1 and S1 (Fig. 1b). From this M1 and S1 were separated into eight and seven clusters, respectively (Fig. 4b). The placement of these clusters along the longitudinal axis was similar to a previous study showing body parts across a somatotopic map (see the subparcellations in Figure 8 of Supplementary Neuroanatomical Results in (Glasser et al., 2016)). The result demonstrates the sensitivity of dMRI for measuring topographic connectivity and the organization of neural populations into a somatotopic map.

### Topographic connectivity over the entire cerebral cortex

We investigated the TF of the entire cerebral cortex to determine how topographic connectivity is spatially organized over the entire cerebral cortex. Fig. 5 shows three paired regions with high, middle, and low TF calculated by Procrustes analysis (Eq. (3)) for subject 100307, where much clearer topographic pattern was visually observed in M1-S1 than V1-V3 and V6-V7. Fig. 6a shows a long tailed distribution of TF for 451 pairs of cortical regions (teal) and shuffled data (gray) by the random rematch of seed voxels and mapping coordinates. The TF distribution (peak: 0.311, median: 0.340, kernel density estimation with Gaussian kernel and Scott’s bandwidth (Scott, 2015)) was significantly higher than the shuffled data (peak: 0.015, median: 0.018, Mann-Whitney U-test, *P* <0.001). Table 1 displays the top 20 paired regions with the highest TF (see Appendix A regarding area name and description). The early sensory areas, including the sensorimotor cortex, visual cortex and auditory cortex, exhibited higher topographic connectivity. Fig. 6c shows that more than 0.5 of high TF appeared in the sensory regions.

**Table 1.** Top 20 paired regions with the highest TF.

| Factor | Area1 | Area2 |
| --- | --- | --- |
| 0.921 | Primary Motor Cortex | Area 3a |
| 0.907 | Area 3a | Area 3b |
| 0.894 | Primary Motor Cortex | Area 1 |
| 0.892 | Primary Motor Cortex | Area 3b |
| 0.873 | Area 1 | Area 3a |
| 0.870 | Area 1 | Area 2 |
| 0.866 | Area 2 | Area 3b |
| 0.865 | Area 1 | Area 3b |
| 0.796 | Third Visual Area | Fourth Visual Area |
| 0.796 | Second Visual Area | Third Visual Area |
| 0.792 | Primary Visual Cortex | Second Visual Area |
| 0.660 | Auditory 4 Complex | Auditory 5 Complex |
| 0.656 | Second Visual Area | Fourth Visual Area |
| 0.629 | Auditory 5 Complex | Area STSd posterior |
| 0.627 | Parieto-Occipital Sulcus Area 2 | Dorsal Transitional Visual Area |
| 0.621 | Primary Motor Cortex | Ventral Area 6 |
| 0.613 | Area 6m anterior | Area 6 anterior |
| 0.593 | Auditory 5 Complex | Area STSd anterior |
| 0.589 | Area OP2-3/VS | Insular Granular Complex |
| 0.583 | ParaBelt Complex | Auditory 4 Complex |

**Figure 5:**
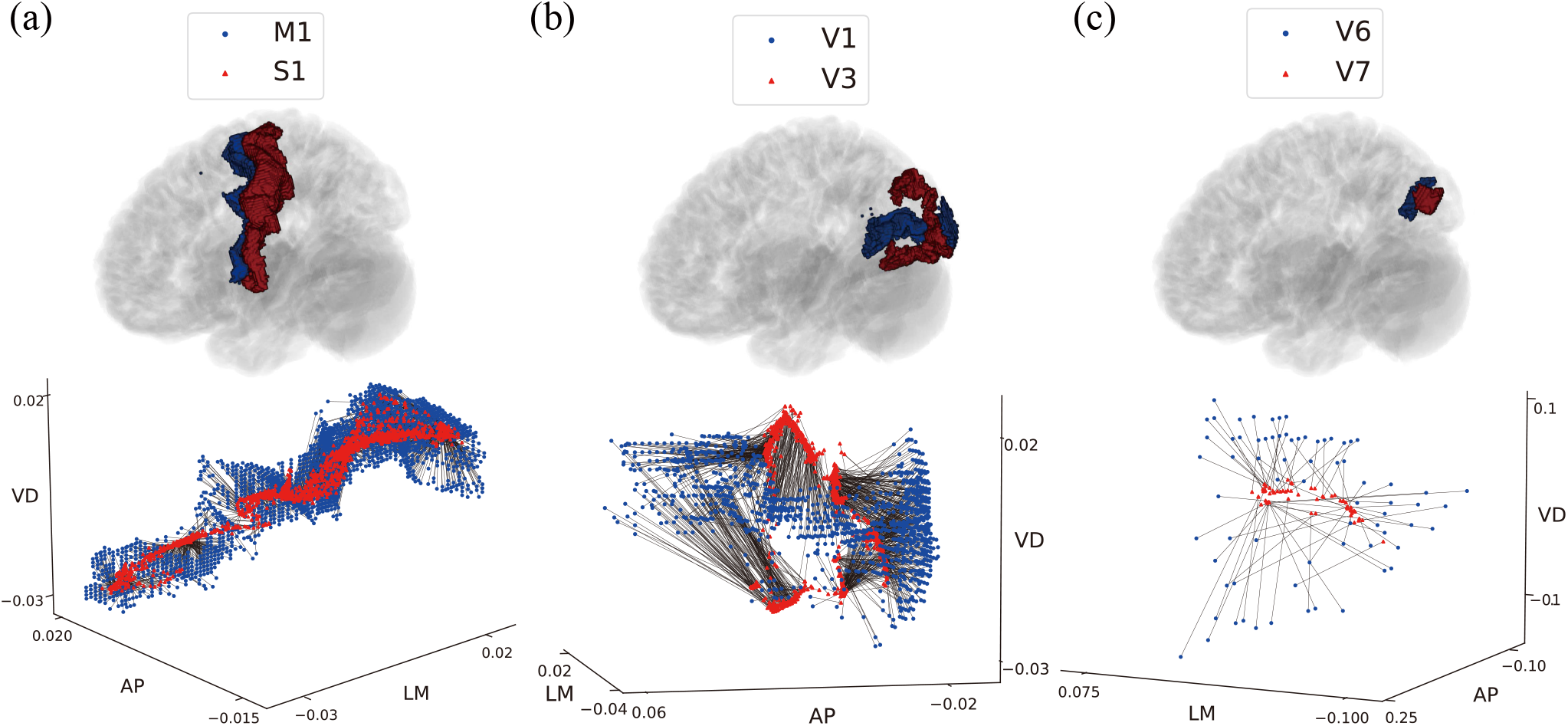
Examples of paired regions to illustrate TF (Subject ID: 100307). (a) Top panel: spatial location of source region M1 (blue) and target region S1 (red) in the left hemisphere. Bottom: connections between seed voxels of M1 (blue points) and mapping coordinates of S1 (red points) after Procrustes analysis, TF=0.880. (b) V1 and V3, TF=0.429. (c) V6 and V7, TF=0.117. (AP: along the anterior to posterior axis; LM: along the lateral to medial axis; VD: along the ventral to dorsal axis.)

**Figure 6:**
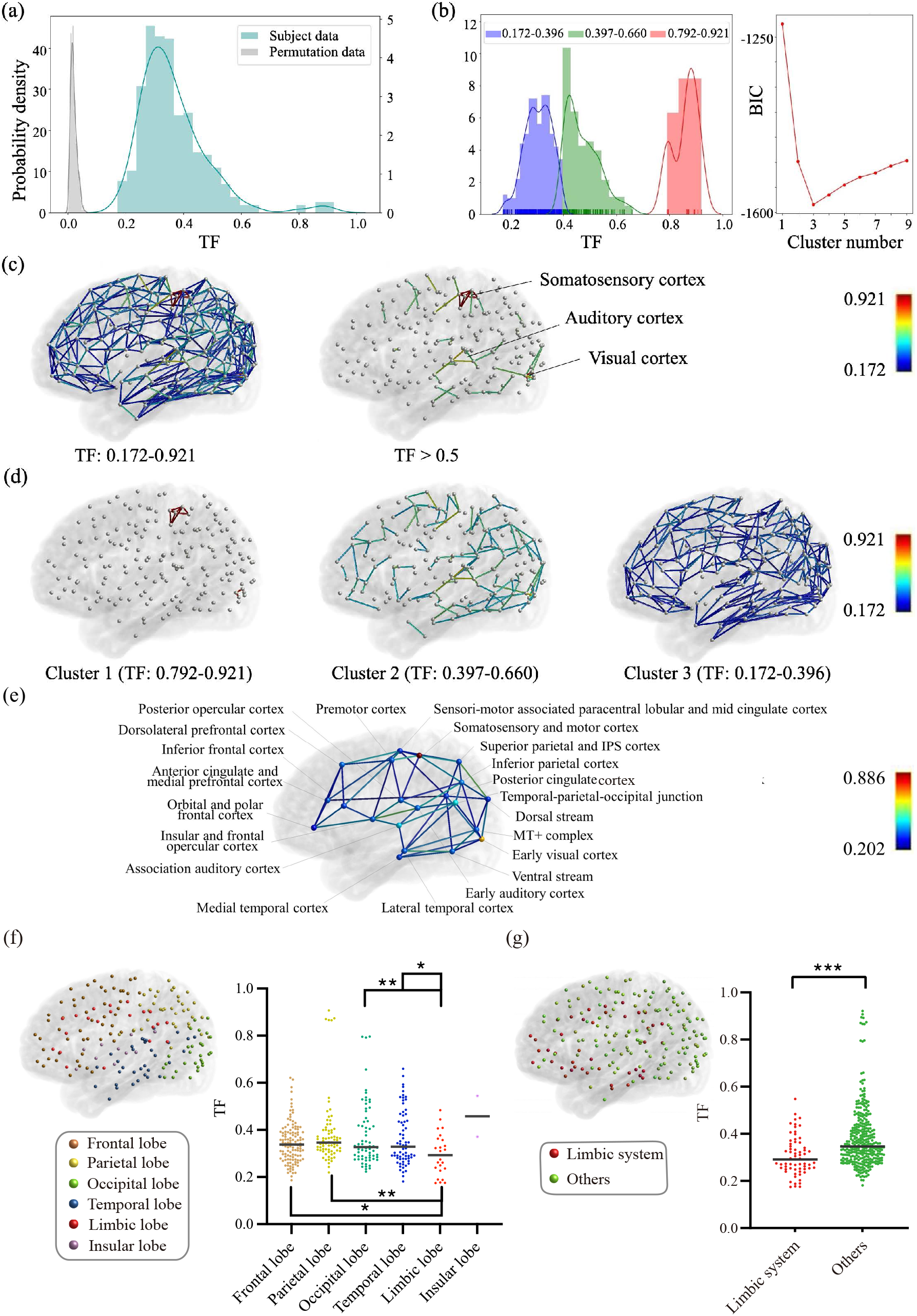
TF over the entire cerebral cortex. (a) Distribution of TF with peak value 0.015 and permutation data with peak value 0.311. (b) GMM clustering were used to generate three clusters of TF. (c) Left: spatial location of TF in cerebral cortex and edge color represents TF value. Right: Edge was drawn when TF surpasses threshold of 0.5. (d) Spatial location of three clusters. (e) Average of TF in intra-group (node) and inter-group (edge). (f) Comparison of intra-lobe TF values among frontal lobe n=120, parietal lobe n=73, occipital lobe n=68, temporal lobe n=69, limbic lobe n=24, and insular lobe n=2, Mann-Whitney U-test. (g) Comparison between the limbic system n=65, and the others n=335, Mann-Whitney U-test. \**P* <0.05, \*\**P* <0.01, and \*\*\**P* <0.001.

Next, we examined the spatial distribution of TFs across the cerebral cortex. We performed GMM clustering for TF values with BIC minima to determine the number of clusters, and then investigated the generated clusters’ spatial locations. The GMM clustering method divided paired ROIs into three subgroups (Fig. 6b, d, and Fig. B1). The first cluster in the right long-tail part of the TF distribution (Fig. 6b red, TF: 0.792-0.921) contained early sensorimotor and visual areas(Fig. 6d cluster 1, Fig. B1 bottom row), which were both located at the distal position of the posterior part from the center of the entire cerebral cortex (Fig. 2). The second cluster (Fig. 6b green, TF: 0.397-0.660) contained primary auditory cortex and sensory areas, which were mainly located around the distal gray matter from the center (Fig. 6d cluster 2, Fig. B1 middle row). The third cluster (Fig. 6b blue, TF: 0.172-0.396) was the most populated and spanned the entirety of cortical regions (Fig. 6d cluster 3, Fig. B1 upper row). These results showed that higher TF regions were distributed around the distal gray matter, especially in the posterior part of the brain. Fig. 6e shows the group average of TF, where 21 groups were defined in HCP_MMP1.0 by geographic proximity and functional similarities (Glasser et al., 2016). The early sensorimotor cortex (0.886) and early visual cortex (0.678) had the highest intra-group TF, while the orbital and polar frontal cortex (0.293), sensorimotor-associated paracentral lobular and mid cingulate cortex (0.323), and posterior cingulate cortex (0.328) had the lowest values.

To check the relationship between anatomical classification and topographic connectivity, we compared TF values among the six cortical lobes: frontal, parietal, occipital, temporal, insular, and limbic lobes (FCAT, 1998) (Fig. 6f, Table A1). The TFs within the frontal, parietal, occipital and temporal lobes were significantly higher than those of the limbic lobe.

Cytoarchitectonic features differ across the cerebral cortex. The limbic system, classified into allocortex, has a 4-layered structure, lower neuronal density, and higher connection density (Catani et al., 2013; Mori and Aggarwal, 2014) (Table A1 and Fig. A1), whereas the isocortex has a 6-layered structure, higher neuronal density, and lower connection density. The limbic system consists of the limbic lobe, orbitofrontal cortex, and anterior insular cortex. To address whether the cytoarchitectonic features reflect topography organization, we performed a statistical test on TF values between the limbic system and the other regions (Fig. 6g). Results showed that the TF within non-limbic system regions were significantly higher than the limbic system (Mann-Whitney U-test, *P* <0.001).

### TF change along hierarchy in visual areas

Previous research using tracer injections reported that the visual area has topographic organization across multiple hierarchical levels (Wang and Burkhalter, 2007). To examine the difference in TF within visual areas, we compared the TF of visual regions from lower to higher hierarchical levels in the ventral (V1->V2->V4->inferior temporal cortex) and dorsal streams (V1->V2->MT->MST) (Perry and Fallah, 2014). The TF in the ventral stream gradually decreased as hierarchical level increased: V1->V2: 0.792; V2->V4: 0.656; V4->PIT: 0.271. In the dorsal stream, although the connection between V2 and MT was not detected in the dMRI data, the TF of MT->MST (0.529) at the higher hierarchical level was less than V1->V2 (0.792) at the lower hierarchical level. These results suggest that topographic connectivity gradually decreases as the hierarchical level increases in visual areas.

### Relationship between TF and spatial features

As shown in Fig. 6, the spatial location of cortical regions, anatomical, and ontogenetic classifications reflect the topographic organization. We further assessed whether the spatial features of regions, such as their shape and size, are critical factors in determining topographic connectivity. To do this, we performed multiple linear regression analysis (MLRA) on the TF values and eight independent features, namely the surface area ratio, minimal surface area, average aspect ratio, ratio of region aspect ratios, average sphericity, ratio of region sphericities, Euclidean distance, and distance from the center of two hemispheres of the cerebral cortex (Fig. 2, see the Methods section for further details). Table 2 summarizes the results of MLRA. The coefficient of determination *R*_2_ was 0.508, and the *F* statistic was less than 0.001. Among these eight spatial features, the minimum surface area (Student’s t-test, *P* <0.001), Euclidean distance (Student’s t-test, *P* <0.001), and average aspect ratio (Student’s t-test, *P* <0.001) were statistically significant factors. The minimum surface area and average aspect ratio showed positive correlations, whereas the Euclidean distance showed an anti-correlation. This suggests that paired regions with high TF tend to be elongated shapes with large surface areas and are closely located. Fig. 7 displays scatter plots between TF and the three significant spatial features, with different colors for different clusters of TF values. The cluster with high TF (Fig. 6d cluster 1) had much larger surface area and aspect ratio than the others, while the spatial features of the other two clusters (Fig. 6d cluster 2, cluster 3) were more closely located.

**Table 2.** Multiple linear regression analysis result between TF and spatial features.

| Spatial feature | Coefficient | t stats | P |
| --- | --- | --- | --- |
| Surface area ratio | 0.105 | 2.007 | 0.045 |
| Minimum surface area | 0.462 | 8.946 | <0.001 |
| Euclidean distance | -0.389 | -10.291 | <0.001 |
| Average aspect ratio | 0.375 | 9.102 | <0.001 |
| The ratio of region aspect ratios | -0.096 | -2.470 | 0.014 |
| Average sphericity | -0.033 | -0.591 | 0.555 |
| The ratio of region sphericities | 0.033 | 0.624 | 0.533 |
| Distance from the center | 0.040 | 0.994 | 0.321 |

**Figure 7:**
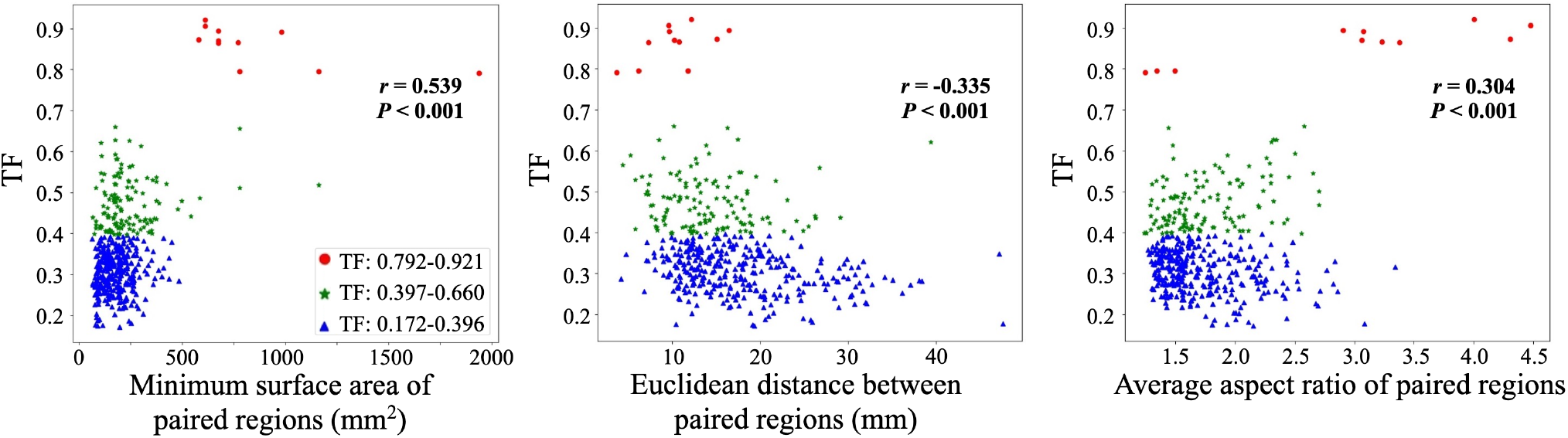
Relationship between TF and three significant spatial features. *r* denotes pearson correlation; points with different colors represent different clusters in Fig. 6b and d.

### Divergent/convergent connectivity

Next, we investigated the divergent/convergent connectivity pattern at two levels: region-to-region, used to evaluate the spatial extent of connectivity between a pair of regions; and global, used to evaluate the spatial extent of connectivity between one region to all other regions in one hemisphere of the cerebral cortex.

### Region-to-region divergence/convergence between a pair of regions

To quantify the degree of divergence/convergence between a pair of regions, we defined region-to-region DC as the average of the standard deviations of terminated target voxel coordinates weighted by the number of streamlines (Eq. (4)). The three highest region-to-region DCs were the pairs dorsal 23 a+b and retrosplenial cortex (RSC), ventral 23 a+b and RSC, and EC and presubiculum (Pres). Fig. 8 displays three examples of high region-to-region DC for subject 100307. Table 3 lists the 20 paired regions with the highest region-to-region DC. Regions with high DC appeared in the medial temporal cortex (RSC and area 23, entorhinal cortex and pres, perirhinal ectorhinal cortex and parahippocampal area 2, lateral parahippocampal cortex and parahippocampal area 2), and anterior cingulate and medial prefrontal cortex (area 33 prime and area 23d, area 10v and area 10r, 33pr and area posterior 24, 10v and orbital frontal complex, 10v and area s32).

**Figure 8:**
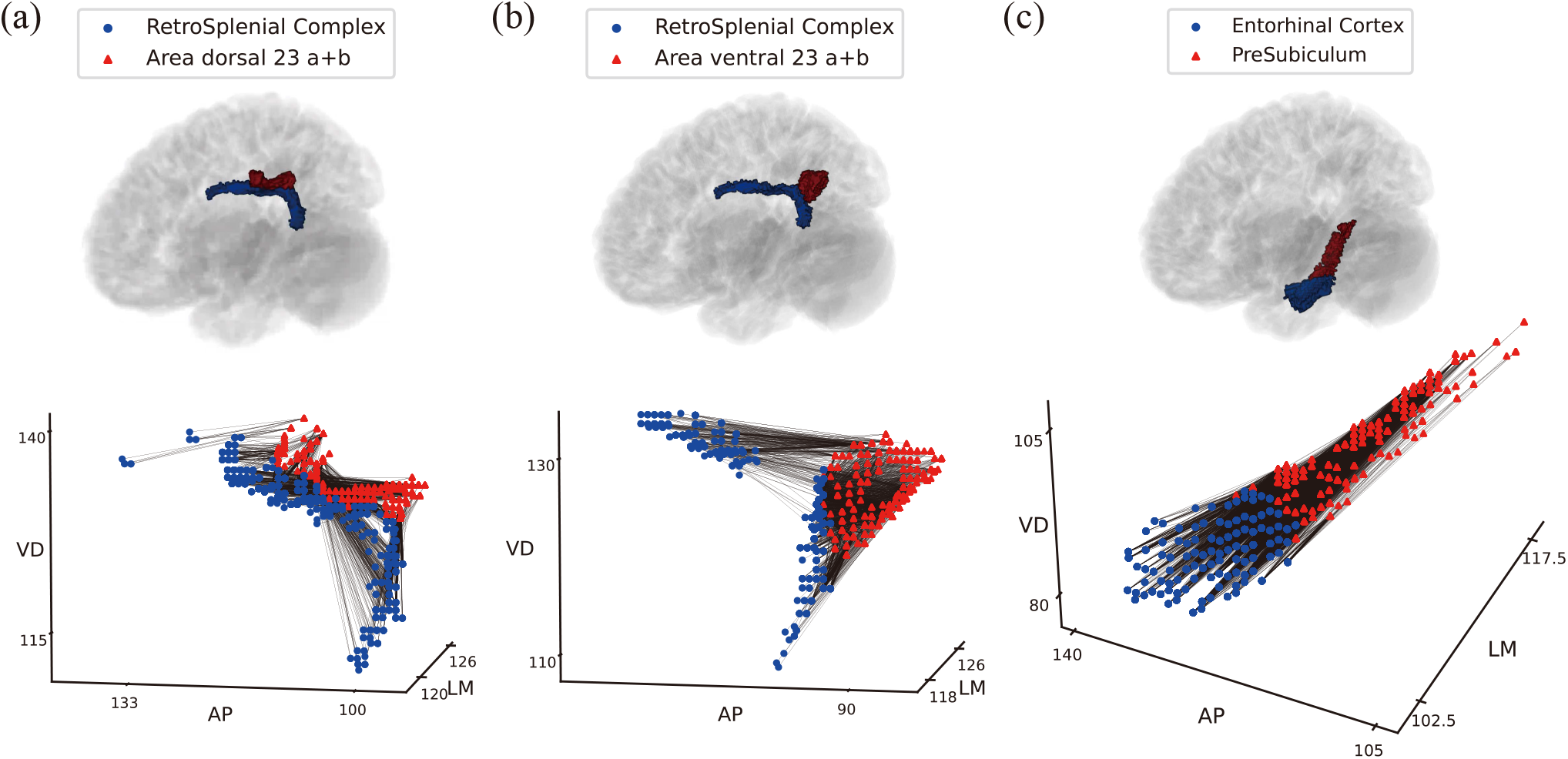
Examples of paired regions to illustrate region-to-region DC. (Subject ID: 100307) (a) Top panel: spatial location of source region (RSC, blue) and target region (Area dorsal 23 a+b, red) in left hemisphere. Bottom panel: connections between seed voxels of RSC (blue points) and target voxels of Area dorsal 23 a+b (red points), region-to-region DC=4.523 and linewidth represents the number of streamline. (b) RSC and area ventral 23 a+b, region-to-region DC=4.569. (c) Entorhinal cortex and presubiculum, region-to-region DC=4.359.

**Table 3.** Top 20 paired regions with the highest region-to-region DC.

| Factor | Area1 | Area2 |
| --- | --- | --- |
| 6.890 | RetroSplenial Complex | Area dorsal 23 a+b |
| 6.646 | RetroSplenial Complex | Area ventral 23 a+b |
| 5.979 | Entorhinal Cortex | PreSubiculum |
| 5.183 | Area 23d | Area 33 prime |
| 5.179 | Area 10v | Area 10r |
| 5.119 | Perirhinal Ectorhinal Cortex | ParaHippocampal Area 2 |
| 4.581 | Area posterior 24 | Area 33 prime |
| 4.563 | Area 10v | Orbital Frontal Complex |
| 4.445 | Area 13l | Orbital Frontal Complex |
| 4.303 | Primary Auditory Cortex | Medial Belt Complex |
| 4.202 | Area 23c | Area Posterior 24 prime |
| 4.174 | Primary Auditory Cortex | Lateral Belt Complex |
| 4.046 | Area 23c | Area 31a |
| 4.016 | VentroMedial Visual Area 3 | Ventral Visual Complex |
| 3.979 | Frontal Opercular Area 3 | Frontal Opercular Area 4 |
| 3.975 | Area TE1 Middle | Area TE2 anterior |
| 3.953 | Area TF | ParaHippocampal Area 3 |
| 3.885 | Auditory 4 Complex | Lateral Belt Complex |
| 3.871 | Primary Motor Cortex | Area 5m |
| 3.867 | Orbital Frontal Cortex | Posterior OFC Complex |

Fig. 9 displays region-to-region DC results for all paired regions in one hemisphere of the cerebral cortex. We performed GMM clustering for the region-to-region DC distribution and examined the relationship between region-to-region DC and spatial features. Region-to-region DC (Fig. 9a teal, peak: 2.502, median: 2.735) had a long tail distribution, which was significantly lower than shuffled data by random rematching of seed voxels and target voxels (Fig. 9a gray, peak: 6.214, median: 6.712, Mann-Whitney U-test, *P* <0.001). Clustering of the region-to-region DC distribution resulted in three clusters (Fig. 9b). Fig. 9c and d show their spatial positions for all clusters and individual clusters, respectively. The first cluster, corresponding to the long tail of the distribution (Fig. 9b red, value: 4.303-6.890), was located in the limbic area (Fig. 9d cluster 1, Fig. B2 bottom row). The second cluster (Fig. 9b green, value: 2.913-4.202) was located around the limbic area, sparsely in the distal gray matter from the center, and neighboring to the pial surface (Fig. 9d cluster 2, Fig. B2 middle row). The third cluster (Fig. 9b blue, value: 1.501-2.912) was the largest and spanned across the entire cerebral cortex, with low density in the limbic area (Fig. 9d cluster 3, Fig. B2 upper row). We also calculated the group average of region-to-region DCs (Fig. 9e). The group with the highest average region-to-region DC was the medial lateral temporal cortex (3.608). The pairs with the top three intergroup region-to-region DC were 1) posterior cingulate cortex and anterior cingulate and medial prefrontal cortex (4.155), 2) early visual cortex and MT+ complex (3.564), and 3) posterior cingulate cortex and sensorimotor-associated paracentral lobular and mid cingulate cortex (3.561).

**Figure 9:**
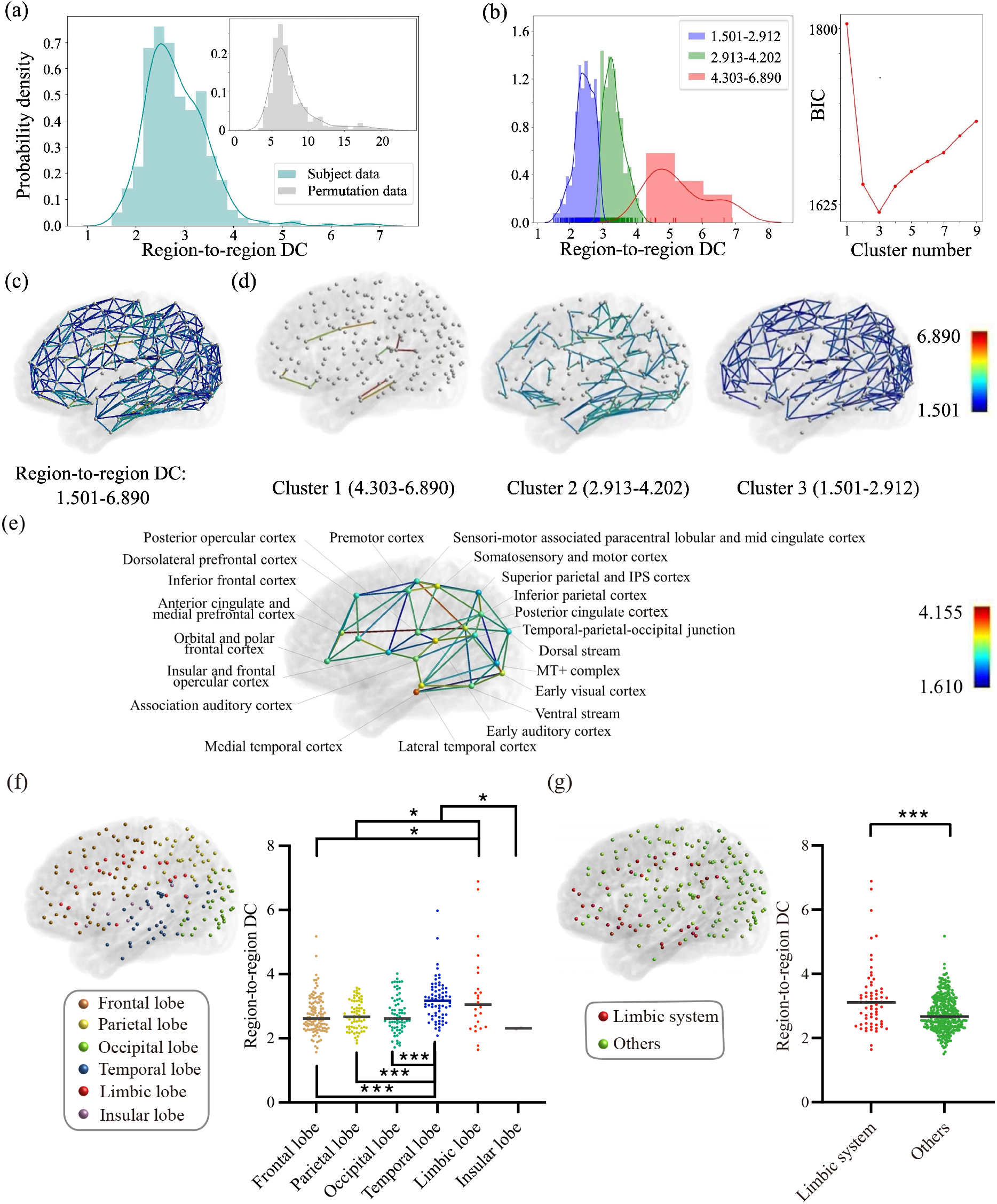
Region-to-region DC over the entire cerebral cortex. (a) Distribution of region-to-region DC with peak value 2.502 and permutation data with peak value 6.214. (b) GMM clustering were used to generate three clusters. (c) Spatial location of region-to-region DC in cerebral cortex. (d) Spatial location of three clusters. (e) Intragroup (node) and intergroup (edge) region-to-region DC. (f) Comparison of intra-lobe region-to-region DC values among frontal lobe n=120, parietal lobe n=73, occipital lobe n=68, temporal lobe n=69, limbic lobe n=24, and insular lobe n=2, Mann-Whitney U-test. (g) Comparison between limbic system n=65, and the others n=335, Mann-Whitney U-test. \**P* <0.05, \*\**P* <0.01, and \*\*\**P* <0.001.

To address the relationship between spatial features and region-to-region DC, we performed MLRA using eight independent spatial features (*R*_2_=0.114, *F* <0.001), as in the analysis of TF. Among these spatial features, the average aspect ratio (Student’s t-test, *P* <0.001, *coefficient*>0) showed a positive correlation, suggesting that paired regions with high region-to-region DC were elongated in shape (Table 4).

**Table 4.** Multiple linear regression analysis result between region-to-region DC and spatial features.

| Spatial feature | Coefficient | t stats | P |
| --- | --- | --- | --- |
| Surface area ratio | 0.055 | 0.781 | 0.435 |
| Minimum surface area | 0.137 | 1.976 | 0.049 |
| Euclidean distance | -0.078 | -1.535 | 0.125 |
| Average aspect ratio | 0.285 | 5.152 | <0.001 |
| The ratio of region aspect ratios | -0.021 | -0.403 | 0.687 |
| Average sphericity | 0.085 | 1.135 | 0.257 |
| The ratio of region sphericities | 0.124 | 1.767 | 0.078 |
| Distance from the center | -0.027 | -0.510 | 0.610 |

We also compared the region-to-region DC among the six cortical lobes defined in the anatomical classification, as for TF. Results showed that the region-to-region DC within the limbic lobe was significantly higher than the others (Fig. 9f). We performed a statistical significance test of region-to-region DC between the limbic system and the other regions (Fig. 9g), showing that the limbic system DCs were significantly higher than the non-limbic system (Mann-Whitney U-test, *P* <0.001).

### Global divergence/convergence between one region and the other regions

To quantify the degree of divergence/convergence connectivity between one region and all other regions in one hemisphere of the cerebral cortex, we defined global DC as the average of the standard deviations of terminated target voxel coordinates weighted by the number of streamlines (Eq. (5)). Fig. 10 shows three examples of source regions with high global DC and Table 5 lists the 20 regions with the highest global DC. Regions with high global DC concentrated around the cingulate and medial temporal cortices in the limbic cortical area (PHA2, 33pr, 23d, 24pr, 23 a+b, p24, RSC, hippocampus, s32, a24, 10v).

**Table 5.** Top 20 regions with the highest global DC.

| Factor | Area |
| --- | --- |
| 10.245 | ParaHippocampal Area 2 |
| 10.180 | Area 33 prime |
| 9.955 | Area 23d |
| 9.530 | Anterior 24 prime |
| 9.530 | Area posterior 24 prime |
| 9.349 | Area dorsal 23 a+b |
| 9.274 | Area posterior 24 |
| 9.230 | RetroSplenial Complex |
| 9.114 | Hippocampus |
| 8.533 | Area 52 |
| 8.493 | Para-Insular Area |
| 8.334 | Entorhinal Cortex |
| 8.235 | Area 23c |
| 8.196 | Sixth Visual Area |
| 8.165 | primary Auditory Cortex |
| 8.150 | Medial Belt Complex |
| 8.146 | Lateral Belt Complex |
| 7.918 | Area s32 |
| 7.890 | Area a24 |
| 7.854 | Area 10v |

**Table 6.** Multiple linear regression analysis result between global DC and spatial features.

| Spatial feature | Coefficient | t stats | P |
| --- | --- | --- | --- |
| Surface area | 0.068 | 0.768 | 0.444 |
| Aspect ratio | 0.372 | 5.546 | <0.001 |
| Sphericity | 0.204 | 2.215 | 0.028 |
| Distance from the center | -0.309 | -4.470 | <0.001 |

**Figure 10:**
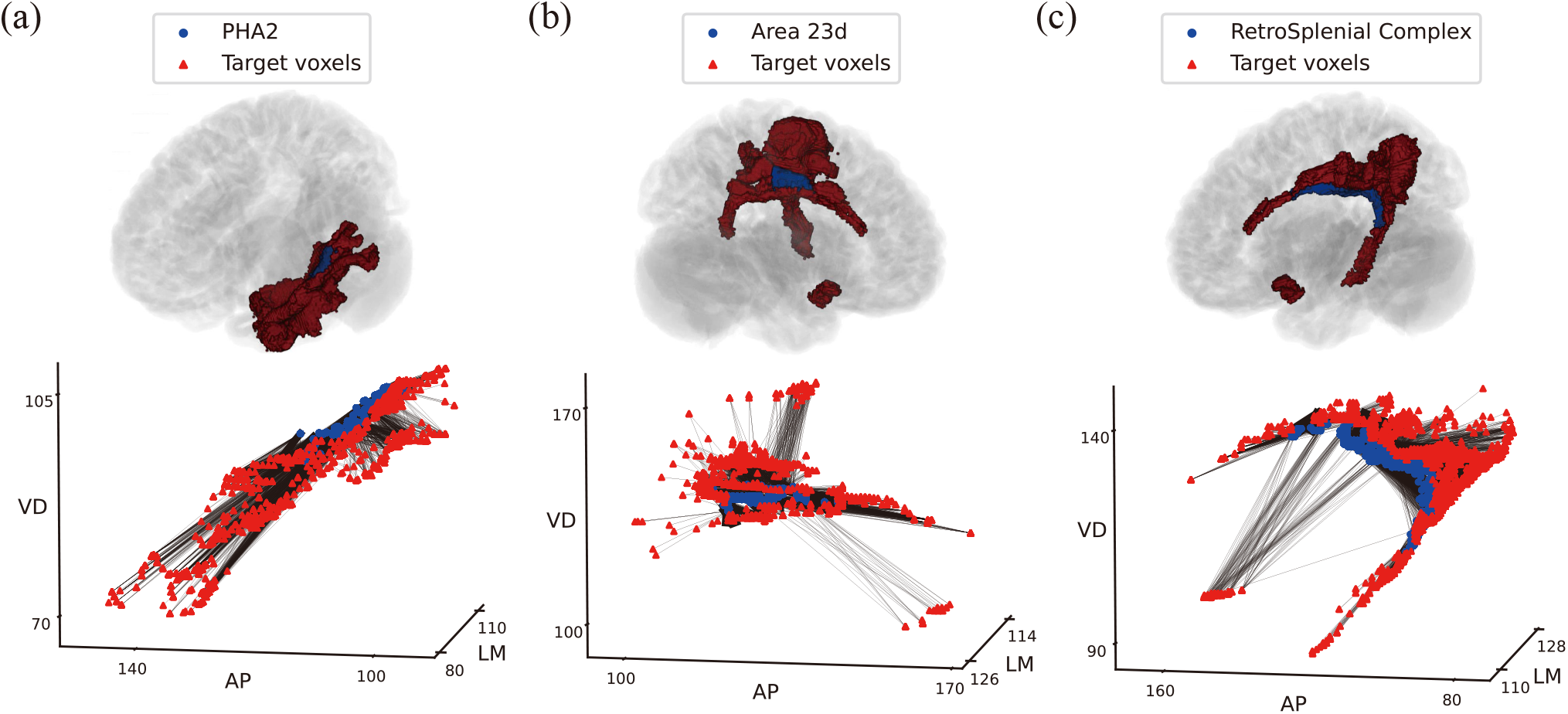
Examples to illustrate global DC (Subject ID: 100307). (a) Top panel: spatial location of source region PHA2 (blue) and target regions. Bottom panel: connections between seed voxels in PHA2 (blue points) and connected target voxels (red points), global DC=10.684 and linewidth represents the number of streamline. (b) Area 23d and connected target voxels, global DC=10.511. (c) RSC and connected target voxels, global DC=9.355.

Fig. 11 displays global DC analysis for all regions in one hemisphere of the cerebral cortex. As for the TF and region-to-region DC analysis, we performed GMM clustering of global DC and examined the relationship between it and a set of spatial features. Global DC (Fig. 11a teal, peak: 5.880, median: 6.181) formed a long tail distribution, which was significantly different from shuffled data, generated by randomly re-selecting the target voxels from all regions in one hemisphere of the cerebral cortex (Fig. 11a gray, peak: 49.422, median: 49.267, Mann-Whitney U-test, *P* <0.001). Clustering of global DC resulted in two clusters; Fig. 11d shows the spatial positions of these two clusters. The first cluster corresponds to the long tail part of the distribution (Fig. 11b, red, global DC:7.193-10.245) and contained regions located in the parahippocampal gyrus, cingulate cortex, and hippocampal formation, which were located at the edge of the medial part and temporal lobe of the brain (Fig. 11d cluster 1, Fig. B3 bottom row). The second cluster was the most populated (Fig. 11b blue, global DC: 3.982-7.097) and contained widespread regions, including distal gray matter from the center (Fig. 11d cluster 2, Fig. B3 upper row). Fig. 11e shows the group average of global DC, indicating high global DC in the medial temporal cortex (7.863), early auditory cortex (7.664), anterior cingulate and medial prefrontal cortex (7.536) and posterior cingulate cortex (7.211).

**Figure 11:**
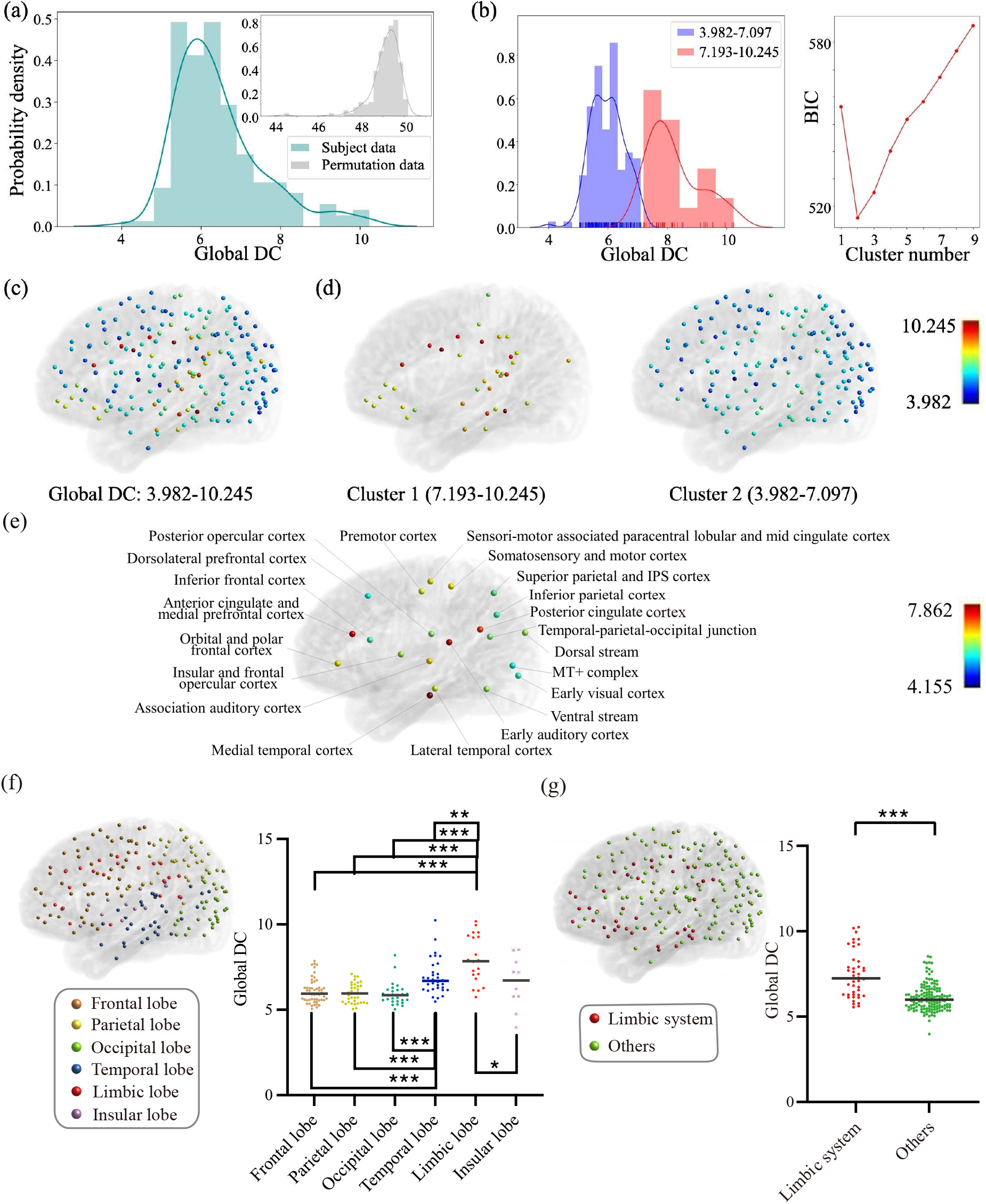
Global DC over the entire cerebral cortex. (a) Distribution of global DC with peak value 5.880 and permutation data with peak value 49.422. (b) GMM clustering were used to generate two clusters. (c) Spatial location of global DC in cerebral cortex. (d) Spatial location of two clusters. (e) Intragroup global DC. (f) Comparison of global DC value among frontal lobe n=52, parietal lobe n=36, occipital lobe n=28, temporal lobe n=33, limbic lobe n=21, and insular lobe n=10, Mann-Whitney U-test. (g) Comparison between limbic system n=42, and the others n=138, Mann-Whitney U-test. \**P* <0.05, \*\**P* <0.01, and \*\*\**P* <0.001.

We also compared global DC among the six cortical lobes and found that the global DC of the limbic lobe was significantly higher than that of the other lobes (Fig. 11f). We performed a statistical significance test of global DC values between the limbic system and the other regions (Fig. 11g), showing that the limbic system DCs were significantly higher than the non-limbic system (Mann-Whitney U-test, *P* <0.001).

To address the relationship between spatial features and global DC, we performed MLRA with four independent spatial features: surface area, aspect ratio, sphericity, and distance from the center. Note that the number of spatial features was smaller than those used for TF and region-to-region DC because the spatial features were only for the source region. Results showed that, to some extent, these features could predict global DC (*R*_2_=0.365, *F* <0.001). The aspect ratio showed a significant positive correlation with global DC (Student’s t-test, *coefficient*>0, *P* <0.001); while distance from the center showed an anti-correlation (Student’s t-test, *coefficient*<0, *P* <0.001), suggesting that regions with high global DC were elongated in shape and located in the proximal gray matter from the center of the two hemispheres of the cerebral cortex.

### Cross-analysis of TF and DC

Fig. 6, Fig. 9, and Fig. 11 showed that high TF appeared in the distal gray matter from the center, while high region-to-region and global DC mainly appeared at the proximal gray matter (limbic and temporal lobes). The distance from the center to each region had no significant correlation with TF, weak anti-correlation with region-to-region DC (pearson correlation coefficient: *r*=-0.156, *P* <0.001), and moderate anti-correlation with global DC (pearson correlation coefficient: *r*=-0.496, *P* <0.001). We further performed statistical test to determine whether the distance from the center in the top cluster (cluster 1) was different from the others. The distance of the top cluster was significantly lesser than that of the second cluster in global DC (Mann-Whitney U-test, global DC: first cluster: 42.177; second cluster: 55.907, *P* <0.001), whereas no significant difference was observed for TF and region-to-region DC. These results suggest that the regions with high global DC were spatially localized around the proximal gray matter from the center.

Next, we directly investigated the relationship among these three connectivity factors. To compare different connectivity factors by node, we converted the factors by edge (TF and region-to-region DC) to factors by node. Here, edge values were distributed to the two participant nodes and then for all nodes the average edge value was used. We also analyzed the node-wise connectivity factors for anatomical and cytoarchitectonic classifications, as shown in Figs. B4, B5, B6, and B7. We confirmed a pattern of significantly lower node-wise TF in the limbic lobes and the limbic system, and a pattern of significantly larger node-wise region-to-region DC in limbic and temporal lobes and limbic system, as the edge-wise TF and edge-wise region-to-region DC. In addition, a weak anti-correlation (pearson correlation coefficient: *r*=-0.199, *P* =0.008) was observed between node-wise TF and global DC (Fig. 12a), and a strong positive correlation (pearson correlation coefficient: *r*=0.647, *P* <0.001) occurred between node-wise region-to-region DC and global DC (Fig. 12b). These results are consistent with the anti-correlation between TF and DC, and similar preference for region-to-region/global DC in anatomically and cytoarchitectonically classified regions(Figs. 9, and 11).

**Figure 12:**
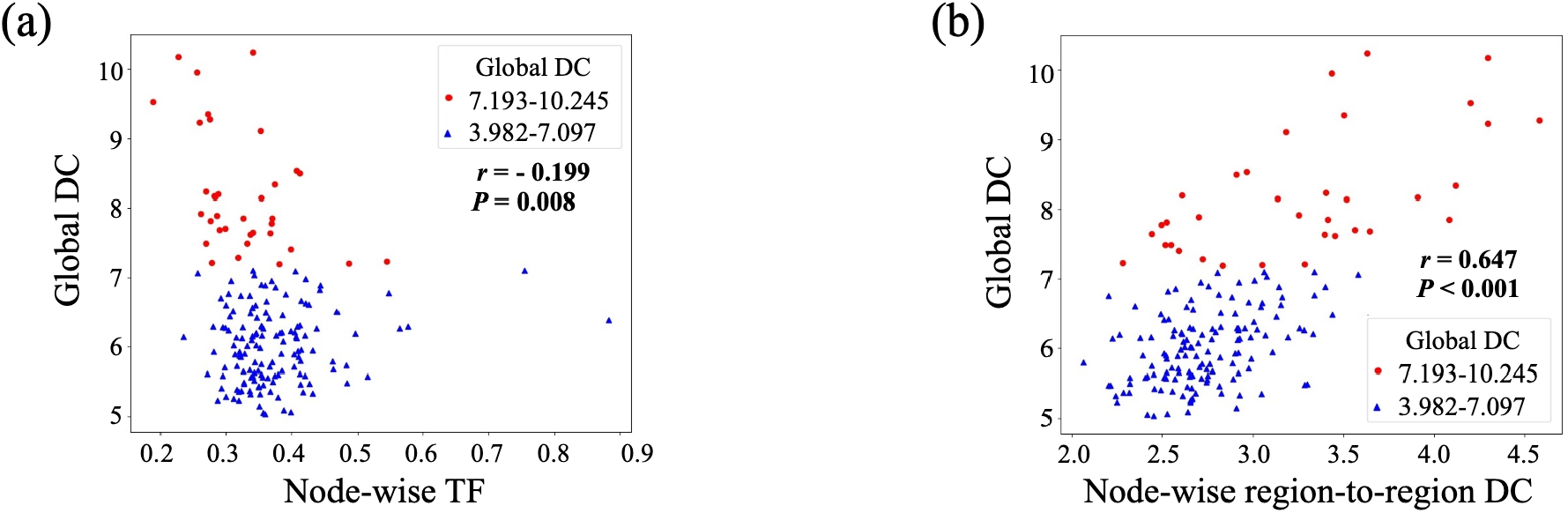
Cross analysis among TF and DC. (a) Scatter plot of node-wise TF and global DC. (b) Scatter plot of node-wise region-to-region DC and global DC. The colors of dots denote two clusters of global DC.

We further checked the degree of overlap of the spatial distribution between region-to-region DC and global DC (Fig. 13). We found that in the limbic system and the top cluster of global DC, region-to-region DC was significantly higher than in other regions (Mann-Whitney U-test, median of region-to-region DC in the limbic system: 3.108; median in the other regions: 2.668; *P* <0.001. median of region-to-region DC in the top cluster: 3.307; median in the other clusters: 2.712; *P* <0.001). These results suggest that regions with region-to-region divergent/convergent connectivity overlaps with the regions with high global DC divergent/convergent connectivity in the limbic system.

**Figure 13:**
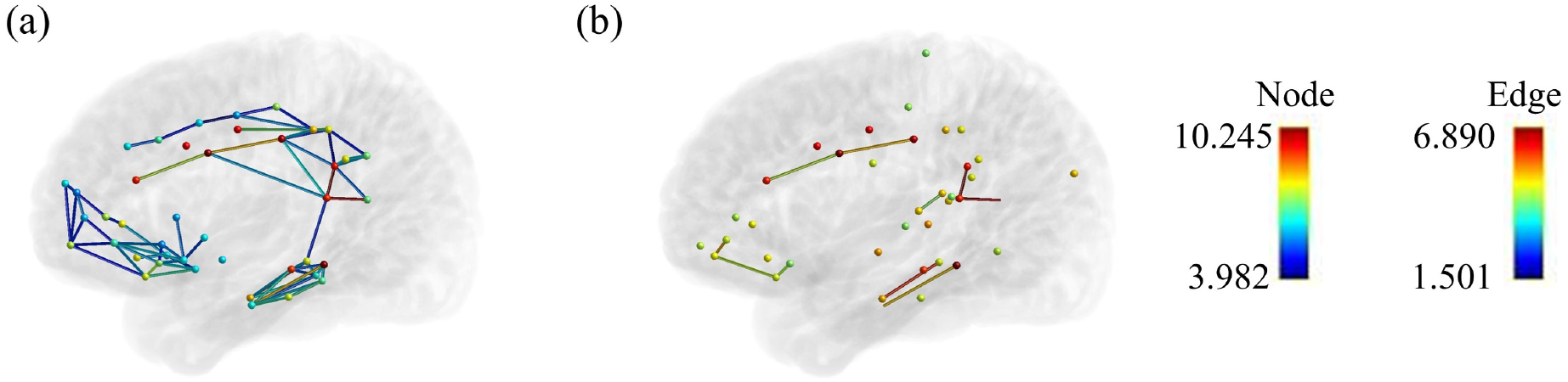
Co-localization of high region-to-region DC and global DC. (a) Superimposition for edges of region-to-region DC and nodes of global DC in the limbic system. (b) Superimposition for edges of region-to-region DC and nodes of global DC in the top cluster of global DC.

## Discussion

Our study analyzed spatial connectivity patterns, topography and divergence/convergence using diffusion MRI data in the human cerebral cortex at the voxel resolution. The summary of the results are as follows: 1) Topographic connectivity, similar to somatotopic mapping, was observed between M1 and S1. 2) The connectivity of M1, S1, and V1 exhibited high TF, whereas the limbic lobe and limbic system showed low TF. 3) TF decreased going from lower to higher order visual area along the ventral and dorsal streams. 4) The temporal and limbic lobes, and the limbic system, showed high region-to-region DC and 5) high global DC. 6) Global DC had anti-correlation with node-wise TF, and a positive correlation with node-wise region-to-region DC.

### Topographic connectivity between M1 and S1

Topographic organization is a typical feature of sensorimotor areas. Previous studies have reported topographic connectivity in the somatotopic map, in the mouse brain using tracer injection data (Zingg et al., 2014) and in the human brain using functional Magnetic Resonance Imaging (fMRI) (Thomas et al., 2021). Our dMRI analysis revealed clear topographic connectivity between M1 and S1 for both manually separated parcels and GMM clustering. In particular, GMM clustering revealed the spatial arrangement of clusters along the longitudinal axis, which is similar to the arrangement of body parts in the somatotopic map previously reported (Glasser et al., 2016). These results demonstrated that the dMRI data used in this study contained essential information on topographic organization and connectivity, reflecting the somatotopic maps in M1 and S1.

### Topographic connectivity over the entire cerebral cortex

Previous research has reported a wide distribution of topographic connectivity in the visual and dorsolateral prefrontal areas of the human brain when using Procrustes analysis with fMRI data (O’Rawe and Leung, 2020). Despite the difference between fMRI and dMRI in spatial resolution and time dimensionality, our results also showed a wide distribution of topographic connectivity, spanning across all cortical lobes. These results support the applicability of dMRI for the comprehensive analysis of topographic connectivity over the entire cerebral cortex.

Topographic connectivity varied among anatomically and cytoarchitectonically classified regions. High TF was observed in early sensorimotor and visual areas (Fig. 6d cluster 1), middle TF in auditory, high-order sensory areas, parietal, premotor, frontal, hippocampal, and temporal areas (Fig. 6d cluster 2), and low TF in the limbic system (Fig. 6g). Although there have been no comprehensive comparisons for anatomical connections over the cerebral cortex, various reports exist on the topographic connectivity and overall organization of the sensorimotor area (Zingg et al., 2014; Glasser et al., 2016; Thomas et al., 2021), visual area (Tootell et al., 1988; Wang and Burkhalter, 2007; O’Rawe and Leung, 2020), the auditory area (Weisz et al., 2004), the olfactory area (O’Leary et al., 1999; Lodovichi et al., 2003), the lateral prefrontal cortex (O’Rawe and Leung, 2020), and the hippocampus and entorhinal cortex (Witter et al., 1993; Tamamaki and Nojyo, 1995; Dolorfo and Amaral, 1998; Jones and Witter, 2007). This universality of topographic features supports the hypothesis that topographic connectivity serves as a general architecture in information processing not only for sensory information in the lower-order cortex, but also cognitive functions, such as analogical reasoning, occurring in the higher-order cortex (Thivierge and Marcus, 2007).

A previous fMRI study using Procrustes analysis showed segregation of unimodal sensory areas and multimodal association areas in gradient decomposition (O’Rawe and Leung, 2020). GMM clustering of TF in the current study separated the primary sensory areas and other higher-order cortical areas containing secondary sensory and multimodal association areas. Although there was a difference in the segregation boundaries between the previous and current study, both separated regions between the low and high hierarchical levels in the cerebral cortex. These results, and the gradual change in TF in the ventral and dorsal streams, suggest that there is a gradient of TF along the hierarchy of levels in the cerebral cortex.

Topographic connectivity has been reported between the cerebral cortex and subcortical regions such as the basal ganglia, cerebellum, and thalamus (Hoover and Strick, 1999; Kishi et al., 2006; Takada et al., 2013; Buckner et al., 2011; Bostan et al., 2013; Sotiropoulos et al., 2016). Moving forward, it would be interesting to examine the relationship between the topographic organization of the cerebral cortex and the subcortical regions in terms of anatomical and cytoarchitectonic classifications.

### Divergent/convergent connectivity over the entire cerebral cortex

We found high region-to-region and global DCs mainly in the medial part of the cerebral cortex, which is largely classified into the limbic system, consisting of the limbic lobe, orbitofrontal cortex, and anterior insular cortex (according to cytoarchitectonic classification (Catani et al., 2013; Mori and Aggarwal, 2014)). The limbic system tends to have high DC for both region-to-region DC, between the different regions that compose it, and global DC, through a spatially wide range of connections over the hemisphere of the cerebral cortex. These results are consistent with previous reports of divergent/convergent connectivity in the limbic system, such as wide afferents of the retrosplenial cortex and posterior cingulate cortex of the macaque monkey (Kobayashi and Amaral, 2003), the global hub structure of the anterior cingulate cortex in the mouse connectome (Coletta et al., 2020), divergent connections in the lateral parahippocampal cortex to the entorhinal cortex (Wellman and Rockland, 1997), the network hub structure of the entorhinal cortex passing convergent cortical inputs toward the hippocampus in the mouse brain (Zingg et al., 2014), and a recipient hub of the entorhinal cortex in the mouse mesoscopic connectome at voxel-level analysis (Coletta et al., 2020).

### Relationship between topographic and divergent /convergent connectivity

Our results showed anti-correlation between node-wise TF and global DC (Fig. 12a). The anti-correlation appeared between limbic and non-limbic systems classified with anatomical and cytoarchitectonic information, where the limbic system had low TF and high DC, and the non-limbic system, containing the early sensory regions, had high TF and low DC. We consider further details of cytoarchitecture and the spatial connectivity organization in the next section. Even the regions with the lowest TF levels in the limbic system with high DC were still significantly higher than the TF of permutation data (Fig. 6), indicating that topographic connectivity and divergent/convergent connectivity coexist rather than being mutually exclusive, despite the anti-correlation. A representative example of this coexistence is in the connectivity between the hippocampus and entorhinal cortex, both of which process episodic memory, spatial mapping, and navigation. Our results showed moderate levels of TF and region-to-region DC between the hippocampus and the entorhinal cortex (Figs. 6, 9). These regions are known to possess both clear topographic and divergent/convergent connectivity (Witter et al., 1989, 1993; Tamamaki and Nojyo, 1995; Dolorfo and Amaral, 1998; Jones and Witter, 2007; Sloviter and Lømo, 2012). The topographic organization and connectivity may work for representing spatio-temporal information (Gu et al., 2018) and the divergent/convergent connectivity may work for associating the information on the spatio-temporal dimensions in the hippocampal system (May-Britt Moser and Moser, 2015).

### Cytoarchitectonic classifications and spatial connectivity

The analysis of spatial connectivity factors with cytoarchitectonic classification showed an anti-correlation between TF and the two types of DC (Figs. 6g, 9g, and 11g). Various cytoacrhitectonic features can explain the difference in spatial connectivity. The primary sensory areas in the posterior part of the brain, belonging to the isocortex, have a six layer structure with granular layer, high cell density (Barbas and Pandya, 1989; Dombrowski et al., 2001; Beul et al., 2017; García-Cabezas et al., 2019; Goulas et al., 2019), and neurons with small dendrites and low spine density (Elston, 2003; Elston et al., 2005). The cytoarchitectonic features of the isocortex fit well with a local computing mechanism through fewer and shorter connections, which is consistent with topographic connectivity (Fig. 6). In contrast, the limbic system, belonging to the allocortex, has a simple layer structure without a granular layer, low cell density, and neurons with large dendrites and high spine density. The cytoarchitectonic features of the allocortex fit with a global computing mechanism through dense and long connections, which is consistent with divergent/convergent connectivity (Figs. 9 and 11). The secondary sensory areas, association areas, and prefrontal areas, which are mainly classified into isocortex, have a medium level of cell and spine densities compared with the primary sensory areas and limbic system. The cytoarchitectonic features of these secondary sensory areas fit as intermediate regions, having high topographic connnectivity with early sensorimotor areas and high divergent/convergent connectivity with the limbic system (discussed in the previous section on the rich-club). Based on the correspondence between cytoarchitectonic features and spatial connectivity, we propose that the cytoarchitecture serves as the basis for determining the balance between topographic and divergent/convergent connectivity.

### Relationship between divergent/convergent connectivity and the rich-club

The limbic system is known as a member of the rich-club of the brain’s network structure (Van Den Heuvel and Sporns, 2011; van den Heuvel and Sporns, 2013). Graph analysis evaluates the topological nature of nodes from the number of connections between nodes irrespective of topographic features and spatial positions. The rich-club is defined as the tendency of nodes connected to many edges (“hubs”), to be very well-connected to each other (Colizza et al., 2006). The major rich-club regions in the cerebral cortex are the limbic system, prefrontal cortex, and parietal cortex (Van Den Heuvel and Sporns, 2011; van den Heuvel and Sporns, 2013). From our results we can interpret that a substantial proportion of connectivity in the rich-club regions is divergent/convergent, and that the regions connecting with them are more spatially distributed. This suggests that rich-club regions may integrate and spread information within itself and with other regions.

The prefrontal and parietal cortex of the rich-club were not classified into the top cluster with high DC in our analysis. That may be attributed to their relative positions in the hierarchy of the cerebral cortex. The prefrontal and parietal cortex communicate with both the sensorimotor areas and the limbic system as the intermediate positions. To communicate with the sensorimotor regions with high topographic connectivity and the limbic system with high divergent/convergent connectivity, the prefrontal and parietal cortex should have both types of connectivities to some degree. To form two types of connectivities, they have the intermediate density of neurons and connections that may cause the medium level of TF and DC as discussed in the previous section. This is the hypothesis of the reason why they were not classified into the first clusters of DC and TF.

### Spatial connectivity organization as economical optimization

The results of this study support the hypothesis that the locations of brain regions and their spatial connectivity properties are optimized for a trade-off between connection cost and information transfer performance (Bullmore and Sporns, 2012). Primary sensory areas with high TF were located distally from the center of two hemispheres (Fig. 6), whereas the limbic system with high region-to-region and global DC was located more proximally (Figs. 9 and 11). In terms of economy of connection construction and signal propagation, topographic connectivity may have high energy efficiency because of shorter connections, while divergent/convergent connectivity may have high information transfer performance because of longer connections. Primary sensory areas in the non-limbic system are interpreted as processing low levels of signals via short connections of topographic connectivity with the large circuity of the distal gray matter. The limbic system, which integrates and broadcasts information, is interpreted to be located at a central position to minimize the total length of connections with the entire cerebral cortex and perform efficient information transfer with divergent/convergent connectivity. Therefore, the current results support the hypothesis of connection cost and information transfer performance being traded off to optimize spatial connectivity and organization in the cerebral cortex. Our results may provide new information on what spatial features can be used for evaluating the balance between connection cost and information transfer performance. It would also be interesting to study how topographic and divergent/convergent connectivities mutually interfere in the developmental process.

### Limitations of the current study

TF is a similarity measure that uses linear transformations, such as shrinkage and rotation, for evaluating the terminated voxel positions’ arrangement between paired regions. When the shape of the source and target region strongly differ in a non-linear way, the capability of the TF is limited. For the precise evaluation of non-linear relationships the development of a new measure is needed.

The sensitivity of TF and DC depends on the voxel size and directionality of connectivity data which dMRI does not have. Spatial connectivity is closely related to the organization of neural population units, such as cortical columns. For example, the horizontal width of orientation columns is less than 1 mm in the visual cortex (Bosking et al., 1997; Kaschube et al., 2010) and topographic organization is observed even within one barrel column in the mouse somatosensory cortex (Andermann and Moore, 2006). The divergent/convergent connectivity among columns representing different information may provide crucial insights into information processing. Also, directed connectivity data can provide information for classifying nodes into sources, that sends outputs, or sinks, that receive inputs (Coletta et al., 2020). Although three types of spatial connectivity were detectable over the entire cerebral cortex in the current analysis using dMRI data with a voxel size of 1.25 × 1.25 × 1.25*mm*, more detailed connectivity information could be obtained with higher resolution data. Furthermore, some important connections observed in tracer injection studies, such as the connection between V2 and MT (Felleman and Essen, 1991), could not be detected in the studied dMRI data, which may originate from limitations in using dMRI data as an analysis approach (Thomas et al., 2014; Calabrese et al., 2015). It is thus important to compare dMRI data with tracer injection data to confirm the limitations of obtaining spatial connectivity information from dMRI data.

The number of ROIs in the brain atlas parcellation also can affect the detection and evaluation of TF and DC. For example, in a past study, a high-resolution parcellation enabled the discovery of rich-club networks in the frontal, parietal, and temporal brain regions (Van Den Heuvel and Sporns, 2011). In the current study, we used the HCP_MMP1.0 parcellation containing 360 cortical ROIs determined from anatomical and multi-modal information (Glasser et al., 2016). Our results demonstrated that this parcellation was sufficient to detect topographic and divergent/convergent connectivity to an extent. It will be necessary to address whether finer-scale parcellation, such as subparcellation representing body parts in the somatotopic map, can be used for obtaining more detailed information.

### Future study

In the modeling of biological spiking neural networks, spatial connectivity information, such as among neurons, layers, columns, and regions, is crucial for building realistic networks (Markram, 2015; Schmidt et al., 2018; Igarashi et al., 2019; Billeh et al., 2020). The results of the current study would be useful for building anatomically detailed models with topographic and divergent/convergent connectivity. The interaction between connectomics and simulation studies may lead to better modeling with a clear critical connectivity pattern.

In this study, we focused on the cerebral cortex of the human brain. Our methods are also applicable to studying different mammalian connectome (SW, 2014; Coletta et al., 2020; Huang et al., 2020), marmoset (Gutierrez et al., 2020; Woodward et al., 2018; Majka, 2020), different brain regions, brain diseases, and brain structural changes after learning. Comparative studies using the measures of topographic and divergent/convergent connectivity would be of interest. Although this study analyzed topography and divergence/convergence as representative connectivity patterns, there are still other patterns, such as reciprocity (Thivierge and Marcus, 2007), and higher-order motifs (Benson et al., 2016; Lotito et al., 2022), that require exploration. Investigation of more complicated connectivity patterns might provide a hint for elucidating the structures for information processing in the brain, although we believe higher resolution connectivity data would be required.

## Conclusion

The current study showed that topographic connectivity was marked in the primary sensory cortices, including the sensorimotor and visual areas, whereas higher region-to-region and global divergent/convergent connectivity was dominant in the limbic system. The contrasting distribution of these connectivity patterns may reflect their functional roles and the trade-off required for the optimization of connection cost and information transfer performance.

## Data availability statement

The data used in this experiment were obtained from a public data source, the WU-Minn Human Connectome Project database (https://db.humanconnectome.org/). All tools used in this study are freely available, including FSL 6.0.4 (https://fsl.fmrib.ox.ac.uk/fsldownloads_registration), ANTs (http://stnava.github.io/ANTs/), Freesurfer 6.0 (https://surfer.nmr.mgh.harvard.edu/), and MRtrix 3 (https://www.mrtrix.org/). Source code (python3.6/shell) used in this study are available upon reasonable request.

## Acknowledgments

This research was supported by MEXT as “Exploratory Challenge on Post-K computer #4-1” (Data-driven modeling of neural networks of the cortex) (Project ID: hp190144 for J. Igarashi), “Program for Promoting Research on the Supercomputer Fugaku” (Project ID: hp200139, hp210169, hp220162 for J. Igarashi), “a Grant-in-Aid for Transformative Research Areas B” (grant number: 21H05137, for J. Igarashi), and the program for Brain Mapping by Integrated Neurotechnologies for Disease Studies (Brain/ MINDS) from the Japan Agency for Medical Research and Development, AMED (grant number: JP15dm0207001 for A. Woodward.). Experimental data were provided by the Human Connectome Project, WU-Minn Consortium (Principal Investigators: David Van Essen and Kamil Ugurbil; 1U54MH091657) funded by the 16 NIH Institutes and Centers that support the NIH Blueprint for Neuroscience Research and by the McDonnell Center for Systems Neuroscience at Washington University. We are grateful to RIKEN’s International Program Associate Program for providing L. Shi the IPA position during this study; Dr. Zhe Sun for helping solve computer issues and constructive suggestions; Dr. Feng Duan for the support of the IPA program; Dr. Ryutaro Himeno, Professor of Juntendo University; and Dr. Satoshi Matusoka, Director of RIKEN Computational Science for their kind support and advice.

## Authorship contribution statement

L. Shi: Conceptualization, methodology, data analysis, writing of original draft, review & editing. A. Woodward: Methodology, review & editing. J. Igarashi: Conceptualization, methodology, writing original draft, review & editing.

## Appendix A Detailed information for 180 ROIs of cortical parcellation in left hemisphere(Glasser et al., 2016)

**Table A1:**
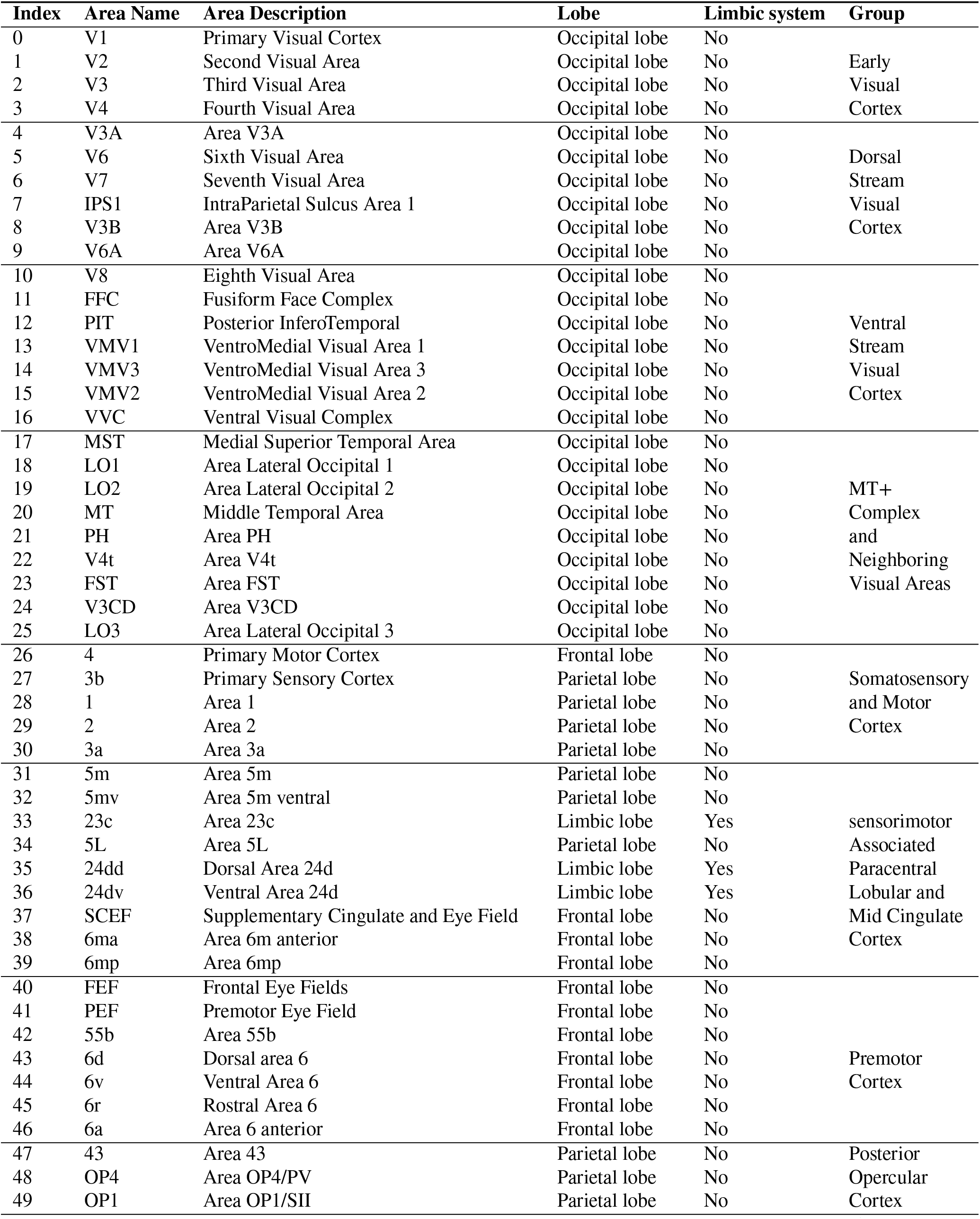
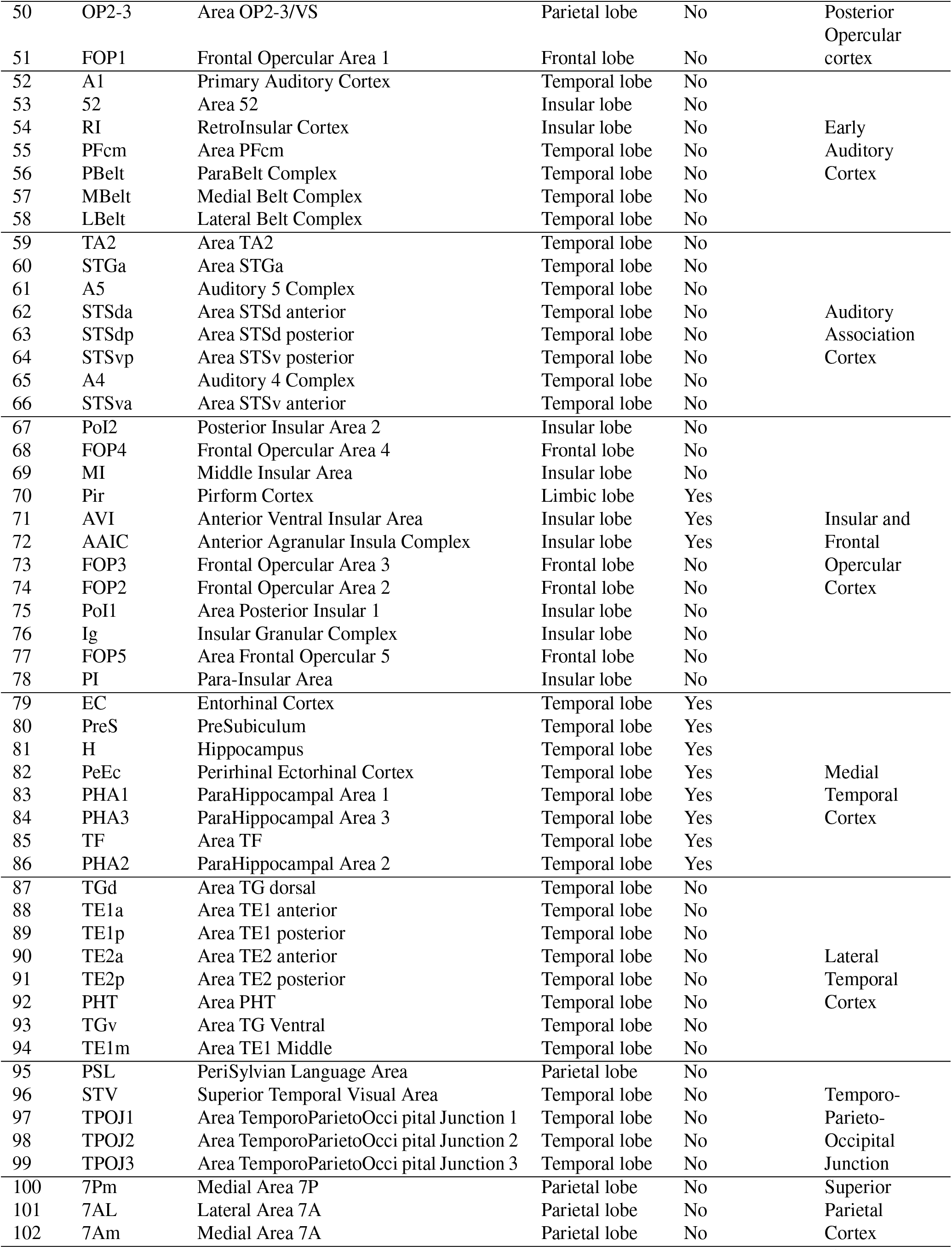
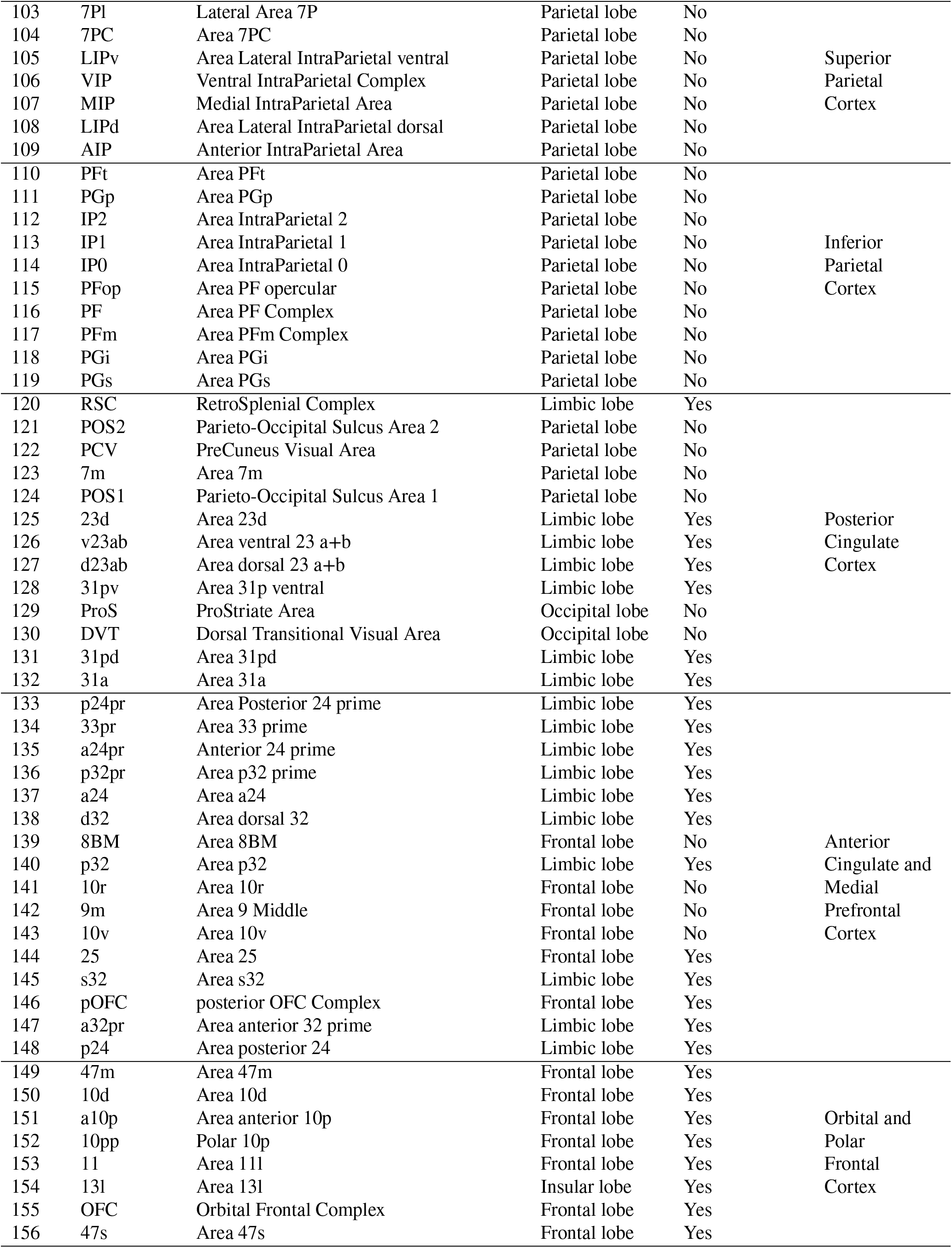
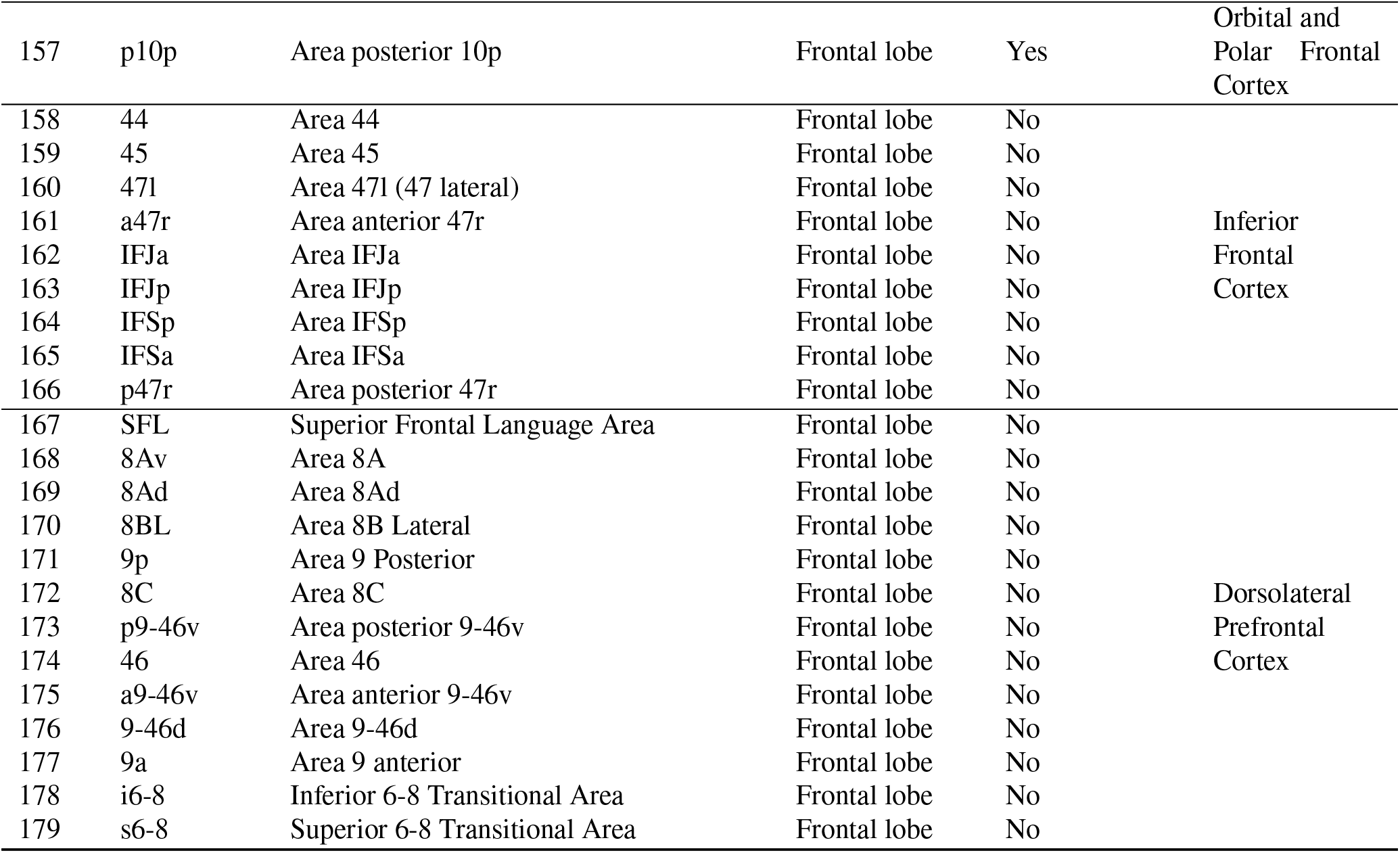
180 ROIs of the cortical parcellation.

**Figure A1:**
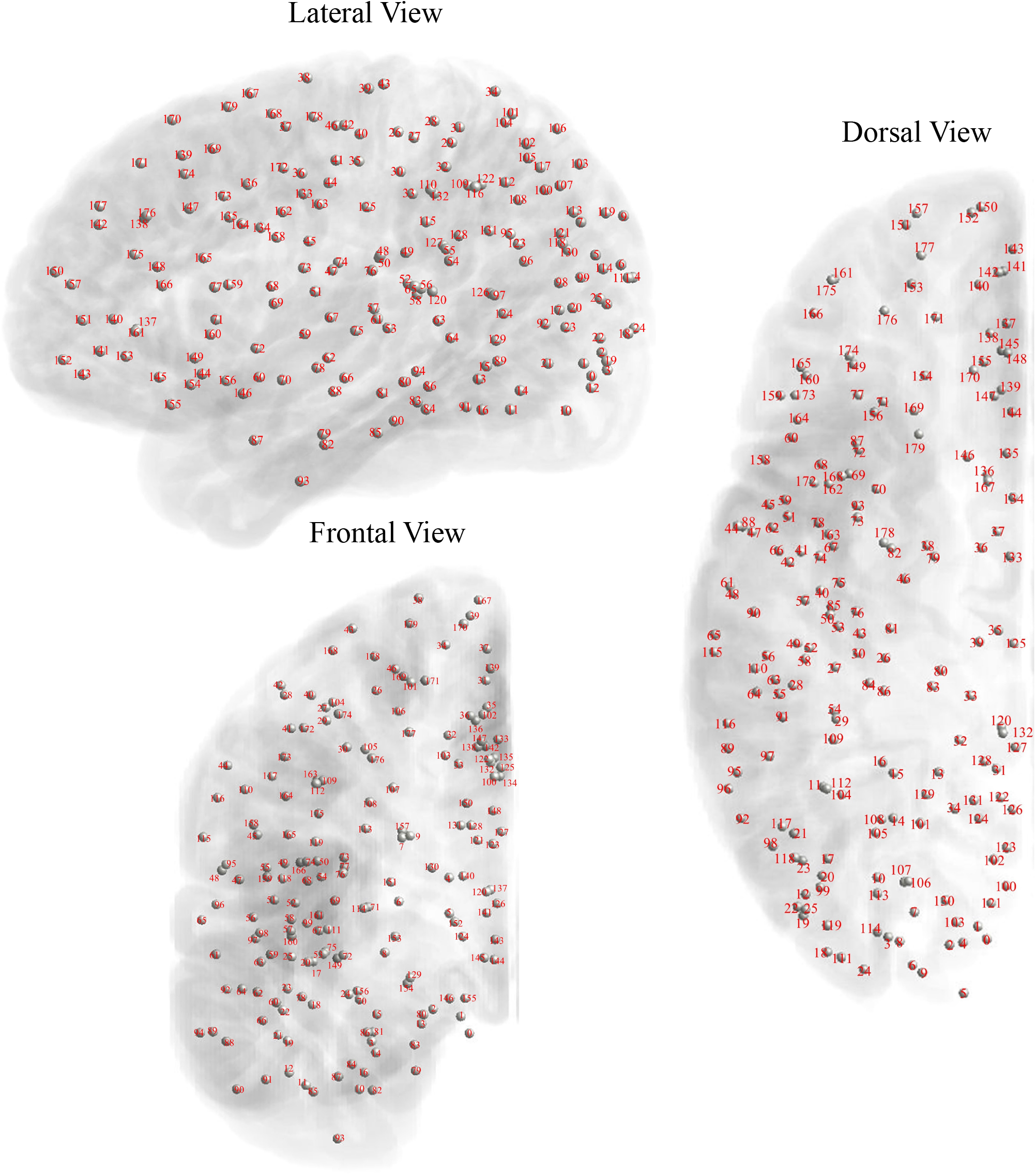
Spatial location and index of 180 ROIs of cortical parcellation in left hemisphere.

## Appendix B Clustering analysis results for TF, region-to-region DC, global DC, node-wise TF, and node-wise region-to-region DC over entire cerebral cortex

**Figure B1:**
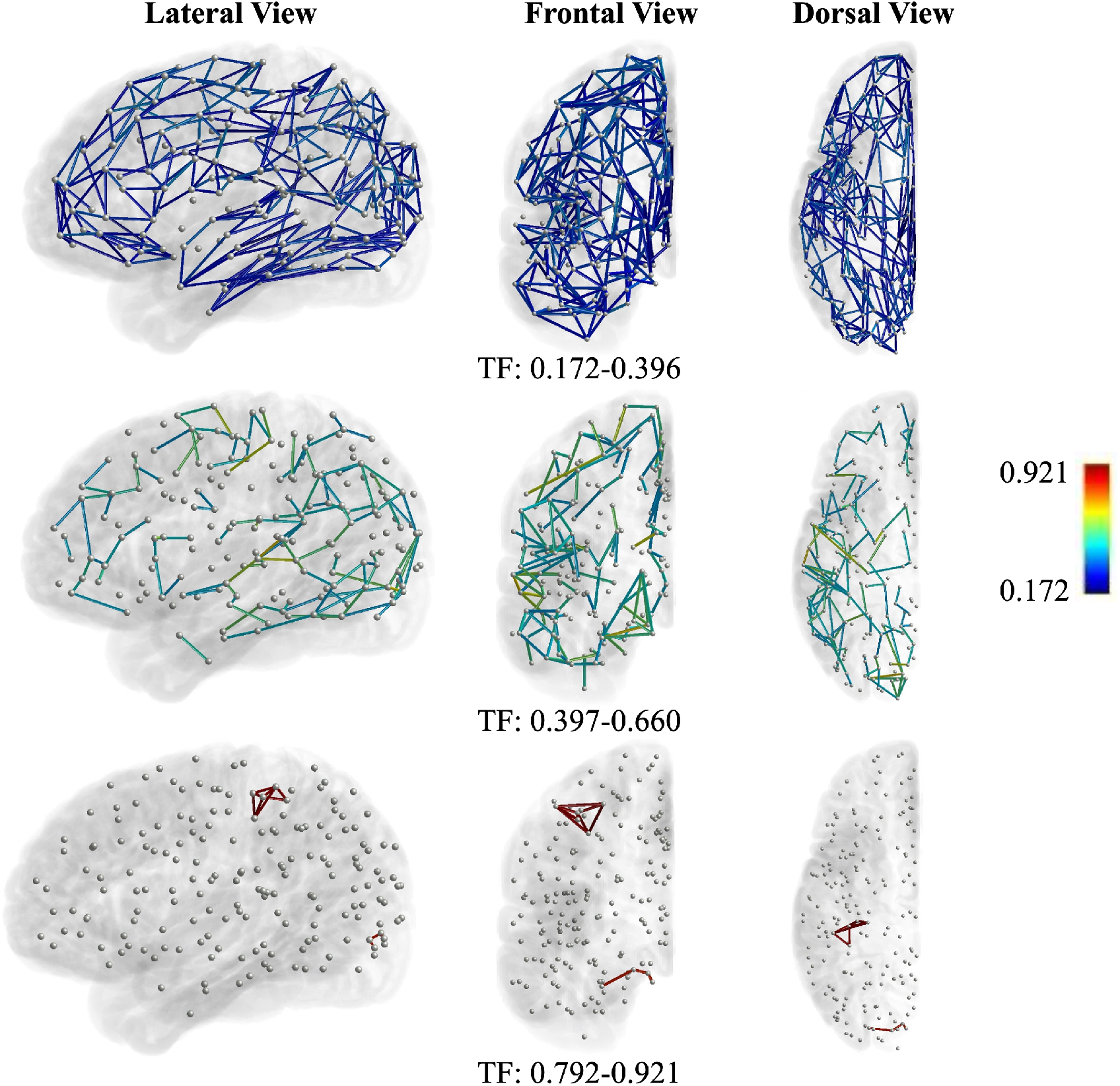
Three clusters of TF in three dimensional space.

**Figure B2:**
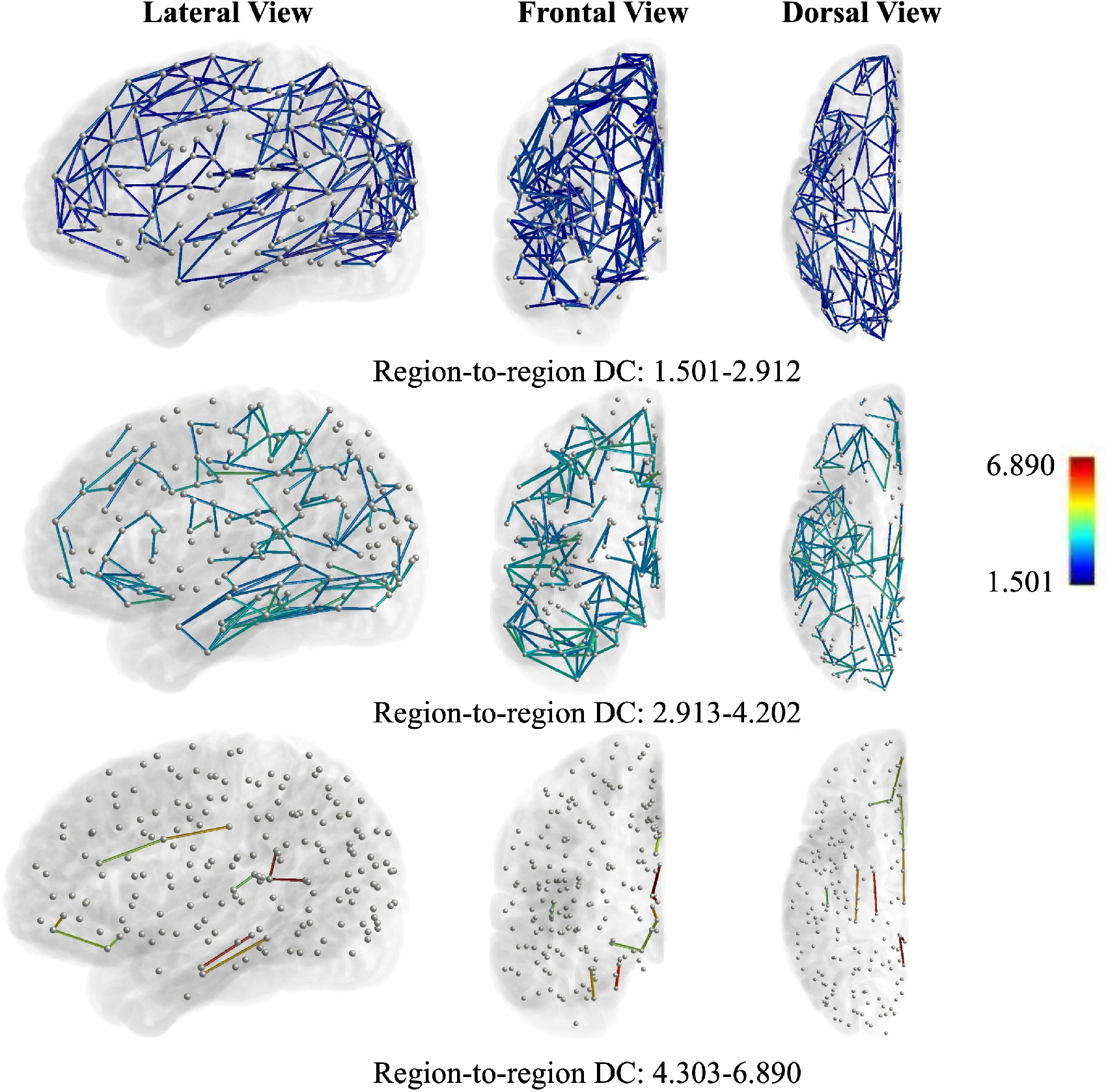
Three clusters of region-to-region DC in three dimensional space.

**Figure B3:**
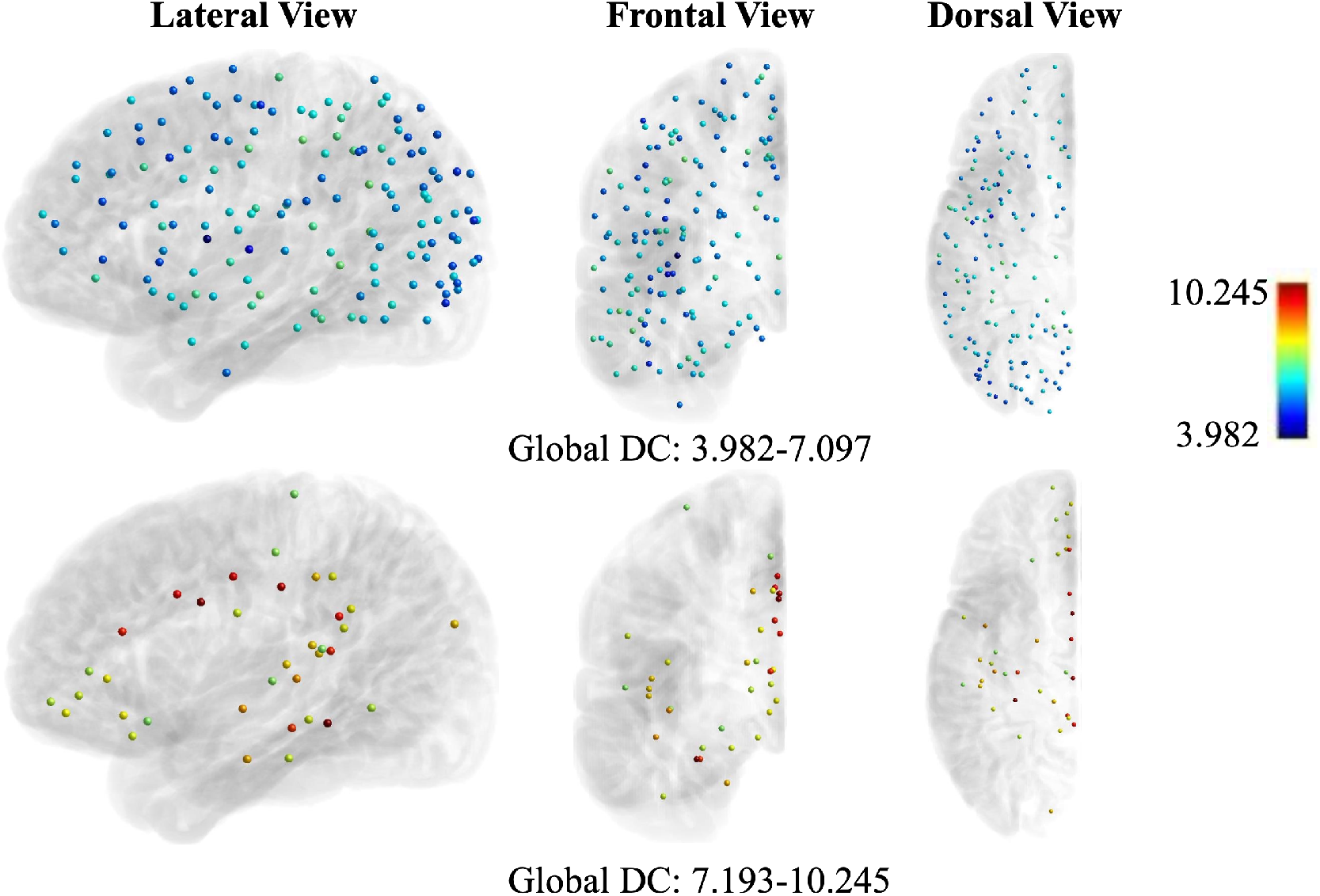
Two clusters of global DC in three dimensional space.

**Figure B4:**
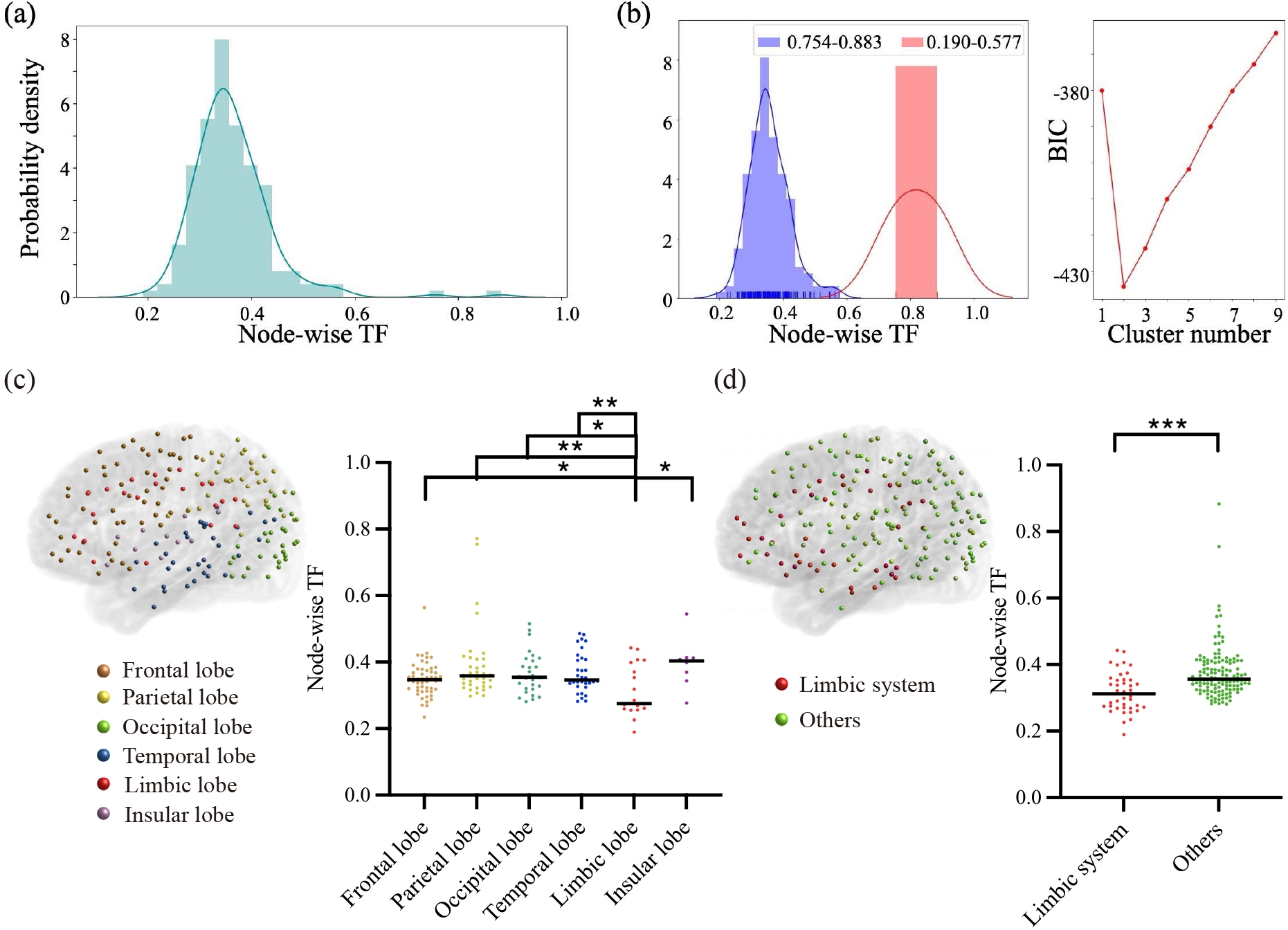
Node-wise TF over the entire cerebral cortex. (a) Distribution of node-wise TF with peak value 0.343. (b) GMM clustering was used to generate two clusters of node-wise TF. (c) Comparison of node-wise TF among frontal lobe n=52, parietal lobe n=36, occipital lobe n=28, temporal lobe n=33, limbic lobe n=19, and insular lobe n=8, Mann-Whitney U-test. Comparison between the limbic system n=40, and the others n=136, Mann-Whitney U-test. \**P* <0.05, \*\**P* <0.01, and \*\*\**P* <0.001.

**Figure B5:**
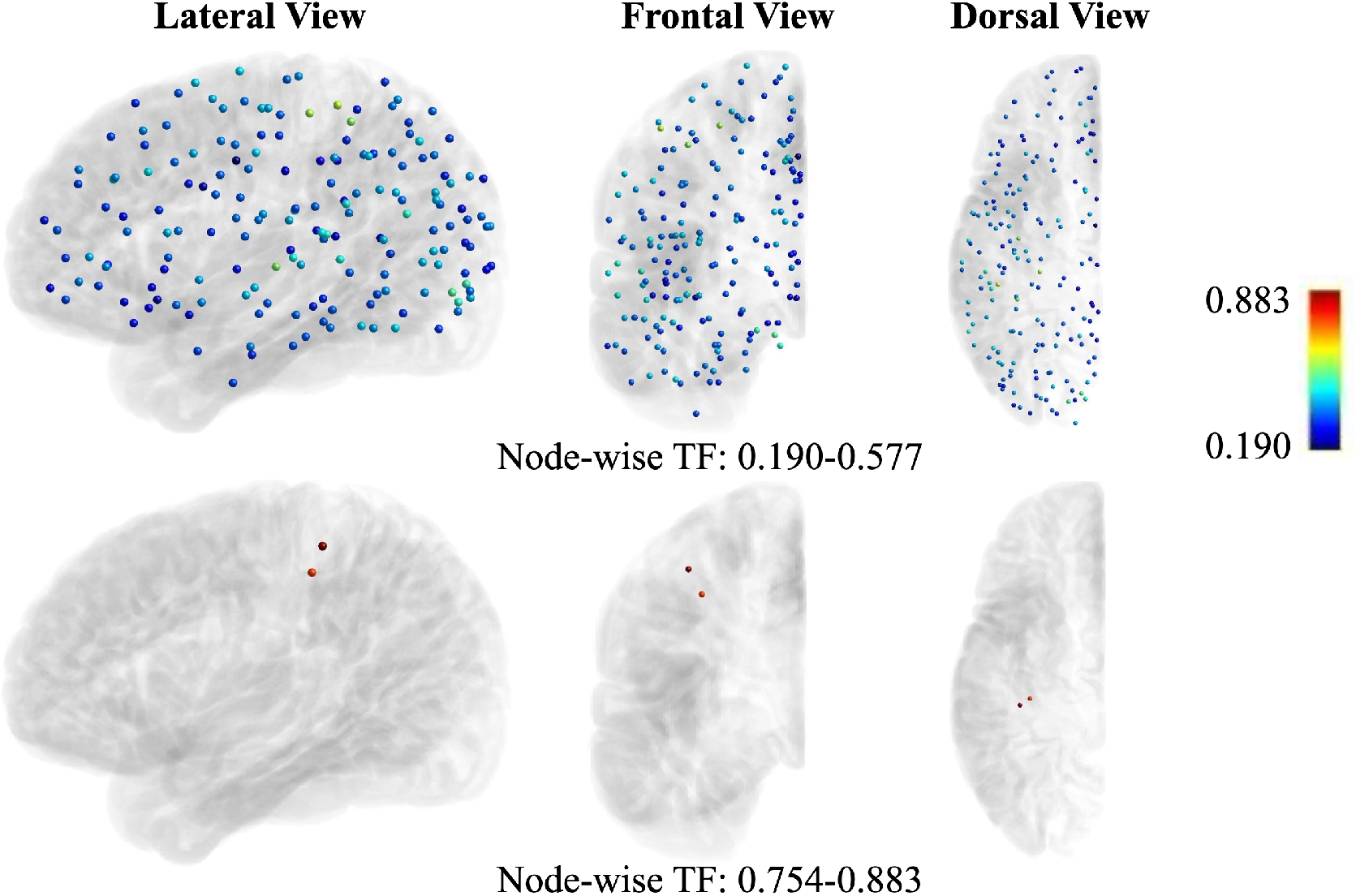
Two clusters of node-wise TF in three dimensional space.

**Figure B6:**
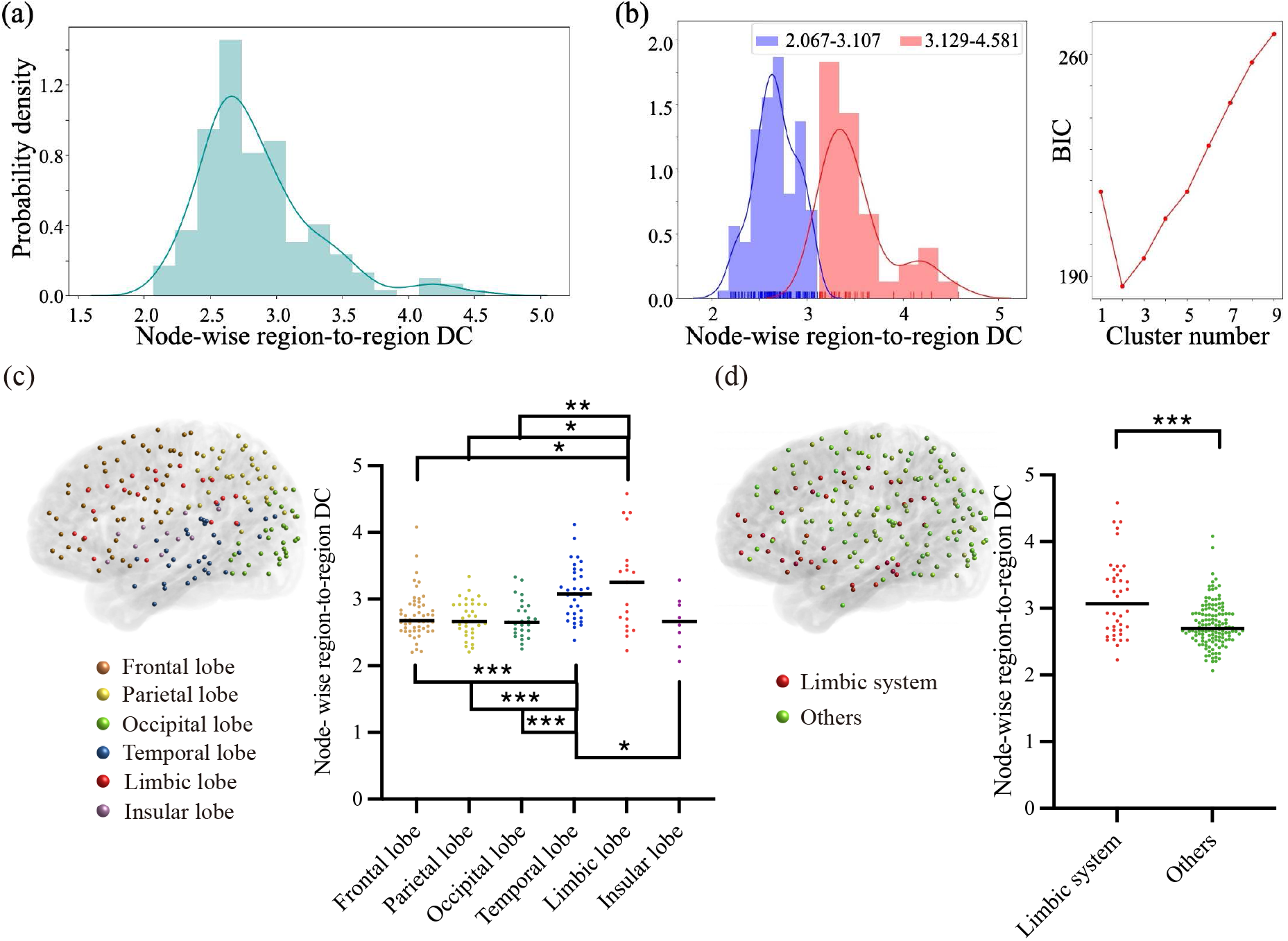
Node-wise region-to-region DC over the entire cerebral cortex. (a) Distribution of node-wise region-to-region DC with peak value 2.657. (b) GMM clustering and BIC were used to generate two clusters. (c) Comparison of node-wise region-to-region DC among frontal lobe n=52, parietal lobe n=36, occipital lobe n=28, temporal lobe n=33, limbic lobe n=19, and insular lobe n=8, Mann-Whitney U-test. (d) Comparison between the limbic system n=40, and the others n=136, Mann-Whitney U-test. \**P* <0.05, \*\**P* <0.01, and \*\*\**P* <0.001.

**Figure B7:**
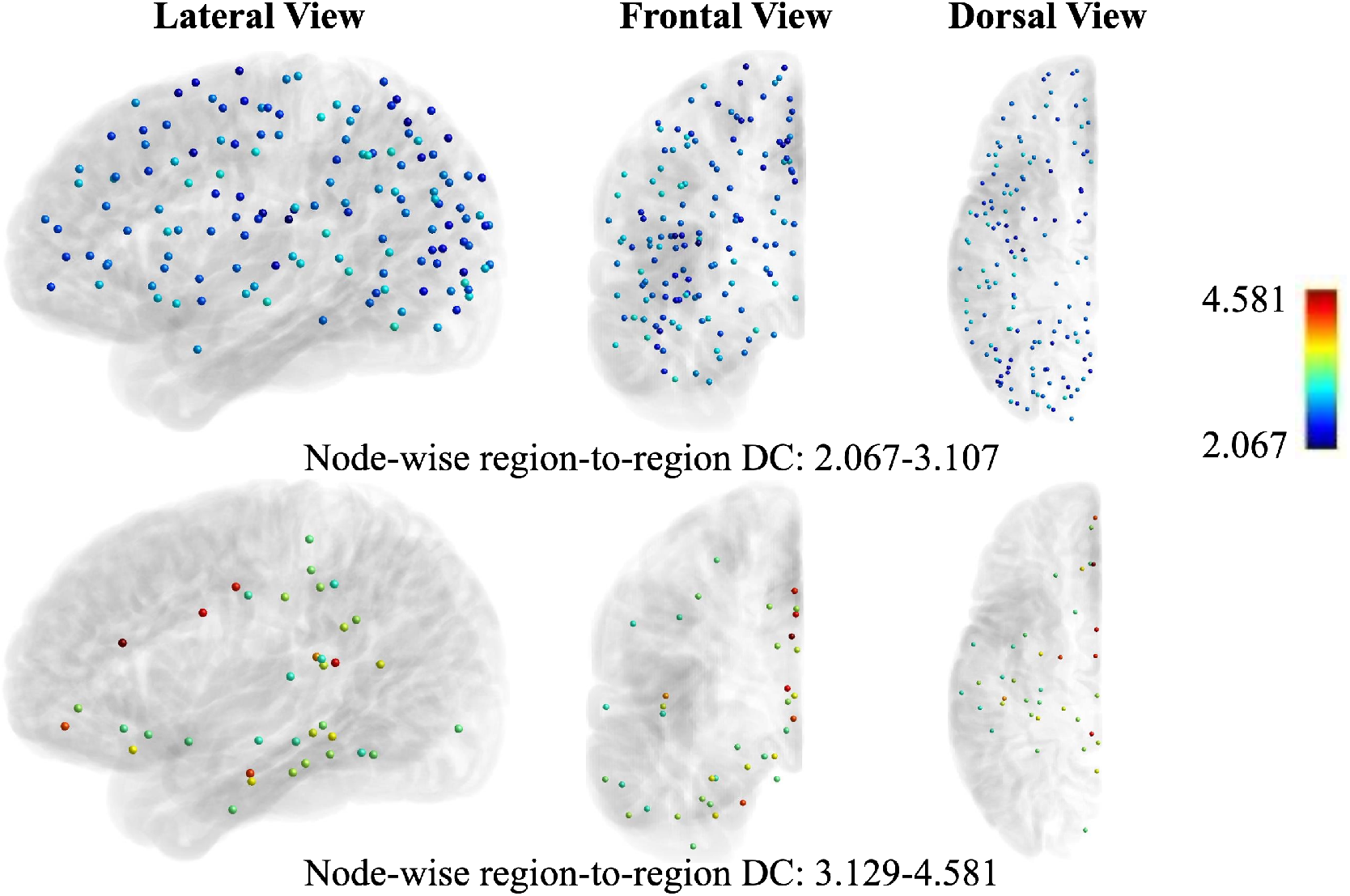
Two clusters of node-wise region-to-region DC in three dimensional space.

## Footnotes

1 https://figshare.com/articles/dataset/HCP-MMP1_0_projected_on_fsaverage/3498446/2

2 https://identifiers.org/neurovault.collection:1549

3 https://github.com/berhane/LAP-solvers

